# *Alcanivorax borkumensis* Biofilms Enhance Oil Degradation By Interfacial Tubulation

**DOI:** 10.1101/2022.08.06.503017

**Authors:** M. Prasad, N. Obana, S.-Z. Lin, K. Sakai, C. Blanch-Mercader, J. Prost, N. Nomura, J.-F. Rupprecht, J. Fattaccioli, A. S. Utada

## Abstract

*Alcanivorax borkumensis* are prominent actors in oil spill bioremediation; however, the interfacial dynamics of their biofilms and its role in oil degradation remain unclear. Longitudinal tracking of biofilm-covered oil microdroplets using microfluidics reveals a spontaneous morphological transition from a thick biofilm phenotype to a thin dendritic phenotype optimized for high oil consumption rates. We show experimentally that biofilm dendrites emerge from aster-like nematic defects in the thin biofilms. We develop a theoretical model that elucidates the transition between phenotypes, linking tubulation to decreased interfacial tension and increased cell hydrophobicity, which we verify experimentally. We demonstrate positional control over the nematic defects on the droplets using microfluidics, causing the biofilm to dimple the droplets. Our results reveal how *A. borkumensis* biofilms utilize topological defects to increase oil access to achieve superior oil consumption rates, which may be a general strategy in oil-consuming bacteria.

**ONE SENTENCE SUMMARY:** *A. borkumensis* adapt their interfacial properties over time to evolve their biofilm phenotype and increase their oil consumption

## INTRODUCTION

Obligately hydrocarbonoclastic bacteria (OHCB) are a group of cosmopolitan marine bacteria with an unusual ecology: they can survive by consuming hydrocarbons as a sole energy and carbon source (*1*). In the marine environment, these bacteria are typically found at very low densities but can bloom to become the dominant bacteria at the site of oil spills (*1, 2*). They are thought to degrade a significant amount of the worldwide spilled oil (*3–5*), which has generated interest in these organisms for their technological potential as agents of bioremediation (*5–8*).

*Alcanivorax borkumensis* SK2 was the first hydrocarbonoclastic bacteria whose genome was sequenced and is frequently used as a model OHCB (*6, 9*). Like most bacteria, it transitions between a free-living lifestyle to one as part of a social collective, called a biofilm, which is now recognized as integral to bacterial biology (*10, 11*). Most biofilms are 3-dimensional communities of densely packed cells that cooperatively secrete extracellular polymeric substances (EPS) that both protect and help the community remain attached to solid surfaces (*12*). This high cell density creates gradients where outward-facing cells have greater access to important nutrients than cells in the interior. However, unlike most biofilm-forming bacteria, *A. borkumensis* form biofilms at the oil-water, a liquid-liquid interface; here, the formation of biofilm creates opposing gradients in the access to oil relative to nutrients needed for respiration. How these nutritive gradients and the fluid nature of the interface affects OHCB biofilm formation and oil consumption remains largely unexplored.

Most knowledge of bacteria-mediated oil degradation comes from genomic sequencing and metagenomic assays (*6, 13, 14*), mutant screens (*15*), interfacial rheology (*16–18*), and microcosm tests (*13, 19, 20*). However, a comprehensive picture of the mechanism of biofilm formation and its coupling to oil degradation remains unclear. To date, few studies have focused on the dynamics of OHCB biofilms formation at the oil-water interface at the micro-drop scale (*21–23*). Nonetheless, these studies do not resolve the physical process of biofilm formation and oil degradation at the single-bacterium scale.

To address these questions, we developed a microfluidic device that allows the trapping and real-time imaging of numerous bacteria-covered oil droplets; this platform allows us to capture the full dynamics of biofilm development starting from individual bacteria through the complete consumption of oil droplets. We show that *A. borkumensis* presents distinct biofilm phenotypes that correlate with culture time using oil. For short culture times, they present as a thick, spherical biofilm (SB) that envelops the droplet. For long culture times, they present as a thin, dendritic biofilm (DB) that stretches and tubulates the drop interface, leading to rapid consumption. We show that tubulation is initiated at aster-like topological defects found in the nematic order of interfacial cells. Using a coarse grain model to describe the biofilm dynamics, we predict that the balance between cell density, interfacial tension, and cell hydrophobicity facilitates a rapid, exponential-like stretching of the interface and confirm this experimentally. Ultimately, the capacity for tubulation at the oil-water interface could be a factor leading to the blooming of *A. borkumensis* after oil spills.

## RESULTS

### Experimental setup and microfluidic device

In pristine marine environments, *A. borkumensis* primarily uses water-soluble organic acids, such as pyruvate, as its carbon and energy source. During marine oil spills, they transition to using water-insoluble alkanes. To study the biofilm dynamics after this transition, we initially cultivate the bacteria using pyruvate, and then switch to a minimal medium supplemented with hexadecane (C16), where we incubate for up to 5 days (d) (**Fig. S1A** and **Supplementary Information (SI) - Methods** section). Using these cultures, we periodically sample the bacteria by generating cell-laden oil microdroplets with fresh C16, which we store in individual traps in our microfluidic platform (*24*). Our device facilitates *in situ* culturing, which enables the longitudinal investigation of biofilm development on specific droplets (**Fig. 1A, Fig. S2A**, and **SI - Microfluidics**). Drops imaged immediately after being trapped initially have ~20-50 cells attached (**Fig. S3A**). These cells divide to form a confluent monolayer over the course of ~12 h. We define the *t*_0_ of our experiments as the moment of confluency.

**Fig. 1:**
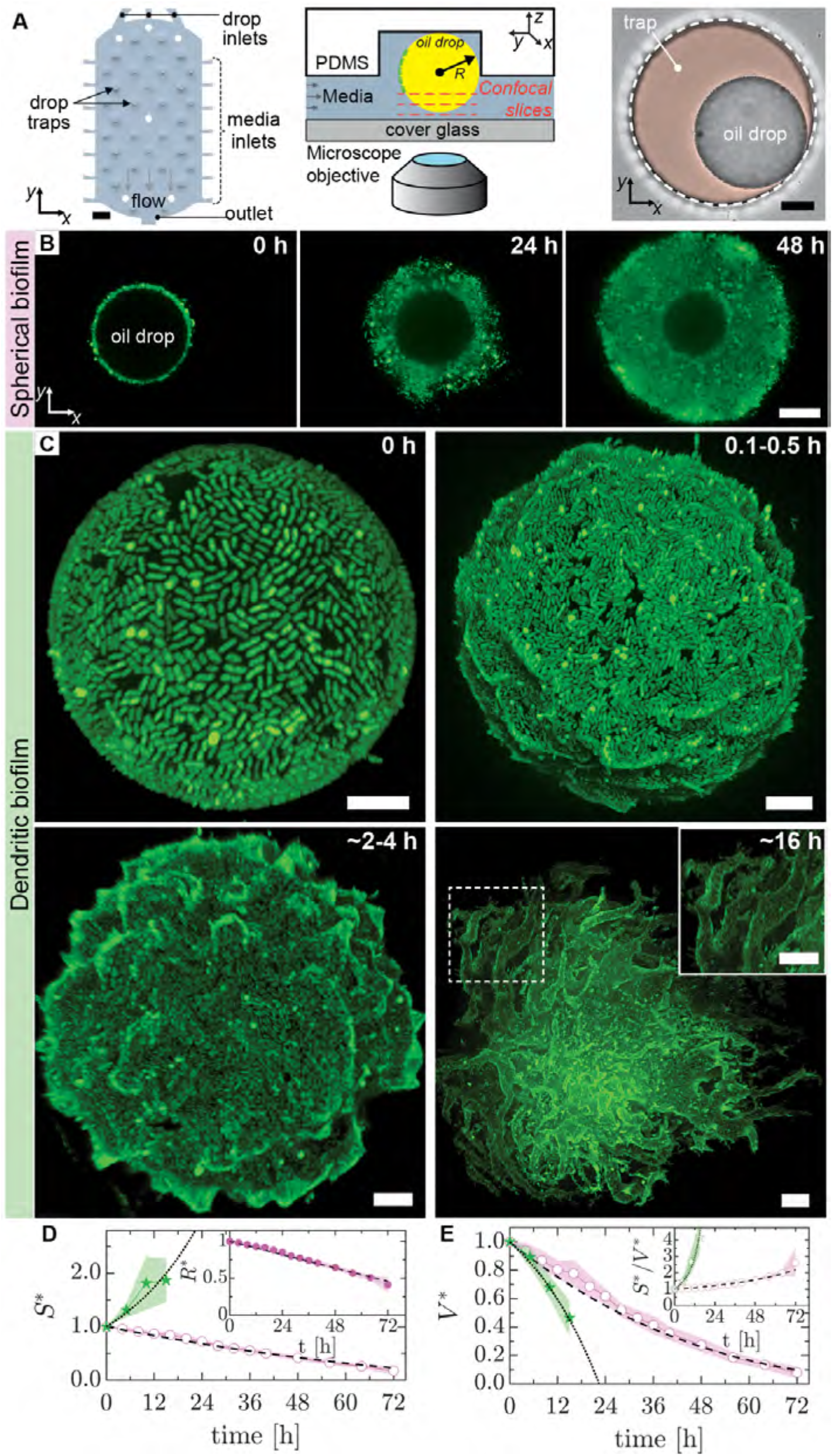
Spherical and dendritic biofilm phenotypes on oil drops in a microfluidic trap. (A) Schematic of the microfluidic oil-drop trap device showing media-filled channels. Oil drops are trapped in the raised circular regions. Inlets on either side of the drop chamber connect to reservoirs that provide a gentle flow of media through the trap chamber. The white circles are pillars. Scale bar =200 *μ*m. (*center*) Schematic cross-section of an individual trap. (*right*) Bright-field image of a representative drop in a trap. The trap is outlined with a white dashed line, while the pocket is colored orange as a guide for the eye. Scale bar = 20 *μ*m. (B) Representative time-lapse sequence of confocal images showing the development of the spherical biofilm (SB) phenotype. As the biofilm (green) grows, the oil droplet (central void) shrinks. Scale bar = 10 *μ*m. (C) Maximum intensity projection confocal images showing the development of the dendritic biofilm (DB) phenotype on representative drops. Local nematic order is present at confluence (~0 h), buckling (0.1-0.5 h) and protrusions (2-4 h) appear later, and large-scale remodeling of the interface leading to the formation of tubes occurs much later (16 h). Scale bar = 10 *μ*m. (D,E) Normalized surface area (*S*\*) and volume (*V*\*) of oil drops as a function of time for (∘) SB and (⋆) DBs. The solid lines and filled regions denote the mean ± one standard deviation (±s.d.) (n_SB_ = 11 drops; n_DB_ = 12 drops). The dashed and dotted lines are best fits of our analytical models of oil degradation for SB and DB phenotypes, respectively (**Supplementary Information**). The filled symbols represent measurements, while open symbols are quantities calculated from measured values. *S*\* for SBs is estimated from the measured normalized drop radius (*R*\*), shown in the inset in (D). *S*\*/*V*\* as a function of time is shown in the inset of (E). *R, S*, and *V* are normalized by their initial values, respectively.

### Bacteria exhibit two distinct phenotypes and consumption rates

Sampling bacteria over 5 d reveals different phenotypes that manifest vastly different biofilm morphologies and oil consumption rates. When sampled after 1 d, *A. borkumensis* form a spherical biofilm (SB) that grows outward from the oil; here, the oil droplet remains spherical as it is consumed **(Fig. 1B, Fig. S3B, and Movie S1)**. In striking contrast, bacteria sampled from 5 d culture can develop into thin biofilms with a local nematic ordering of the interfacial cells, which eventually buckle and then tubulate the droplet interface, as shown in **Fig. 1C**; we call these dendritic biofilms (DB). In this case, the biofilm buckles the interface to accommodate a population that is continuously increasing in number. The magnitude of the deformations grows as the oil is consumed, which ultimately shreds the droplets into tiny fragments (**Fig. S3C and Movie S2)**.

We quantify the oil consumption rate for each phenotype by measuring the droplet volume (*V*) from brightfield and confocal images of the droplet surface area (*S*) over time (**SI - Methods**). SBs consume >90% of the initial volume in ~72 h, whereas DBs achieve the same in ~20 h. We find that the oil volume of SBs decreases as a polynomial function of time, whereas for DBs, the decay is much faster (**Fig. 1D,E** and **SI - Eq.11**). In both cases, the decrease is consistent with a model of oil consumption effected exclusively by bacteria at the interface (**SI - Analytical Model of Oil Consumption**).

The good agreement between our analytical models and data allows us to estimate the per-cell consumption rates of both SB and DB cells as 0.7 and 0.8 fL/h, respectively; this small difference in consumption rate is consistent with their similar division times (**Fig. S1D**). For comparison, the volume of a single cell is ~1 fL, meaning that these bacteria consume a volume of oil close to their own, every hour. Despite the similarity in consumption rates on a per-cell basis, the normalized surface-to-volume ratio (*S*\*/*V*\*) shows that DBs are significantly more efficient: *S*\*/*V*\* doubles in 72 h for SBs, whereas it diverges in less than 24 h for DBs (**Fig. 1E, inset**). The *S*\*/*V*\* ratio provides a means for comparing the relative efficiencies of the two phenotypes and highlights the fact that these differences arise from the rapid increase in interfacial area caused by DB biofilms. In both cases, the shape of the interface defines the dynamics of volume decrease. For SBs, the interfacial area defined by the spherical droplet determines the number of cells (*N*) that can pack onto the interface to have access to the oil. Conversely, for DBs, it is the increase of *N* at the interface due to cell division that determines the interfacial area. Thus, the rate of consumption decreases over time for SBs, while it increases continuously for DBs.

### Tubulation is facilitated by topological defects

We correlate the onset of the rapid increase in surface area for DBs to the emergence of nematic order of the interfacial cells. Numerous examples have shown that rod-like bacteria can locally align in the same direction, defining a nematic order field (*25–28*). Active systems with nematic order have demonstrated the ability, both in theory (*29, 30*) and in experiments (*31–34*), to utilize topological defects to achieve surface deformation. Here, at 2-4 h post-confluency, we observe the appearance of conical protrusions of cells that originate from the core of aster-like (+1) topological defects within the nematic field (**Fig. 2A,B** and **Fig. S4A,B)**. As the biofilm matures, more protrusions appear while existing protrusions elongate into branched bacteria-covered tubes (**Fig. 1C (16 h)** and **Fig. 2C)**.

**Fig. 2:**
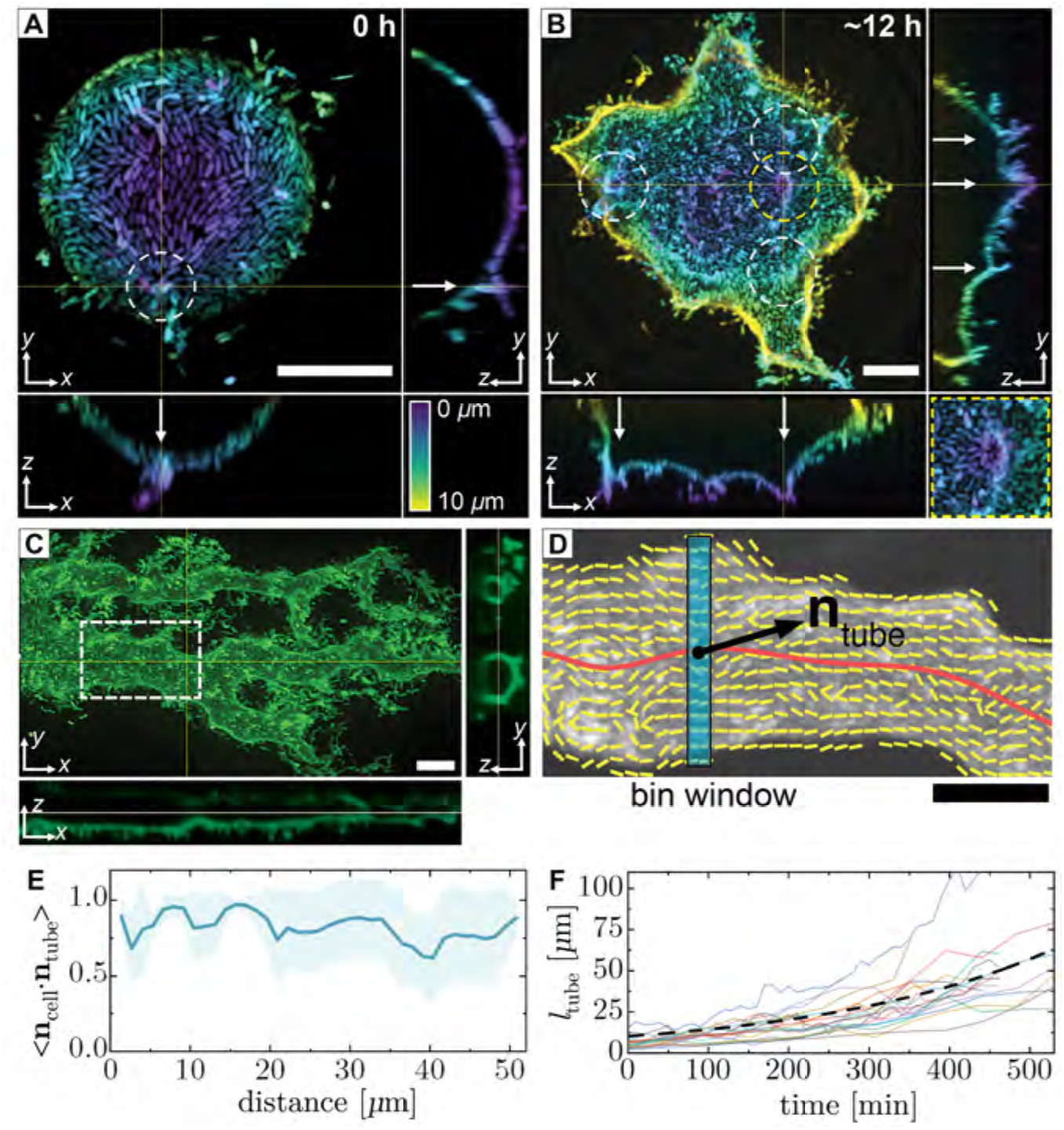
Dendrites originate from topological defects. Confocal images of representative droplets (A) early (~0 h) and (B) later (~12 h) in biofilm development. The images are color coded by depth. The dashed circles enclose topological defects with a charge of +1, while the arrows indicate protrusions. (B, inset) A magnified view of the central defect circled in yellow. (C) Confocal image of a bacteria-covered tube connected to a deformed droplet with corresponding orthogonal views (~20 h). (D) Director field of the visible cells in the dashed box in (C). The director field (yellow lines), tube axis (red line), the local tangent unit vector (**n**_tube_) along the tube axis, and the bin window (blue) are shown. (E) Axial order of the cells along the oil tube shown in (D). For a sliding window of width 1.5 *μ*m along the tube, the local axial order is defined as the average of the individual scalar products between the director-field unit vectors (**n**_cell_) and local tangent unit vectors. <**n**_cell_ ⋅**n**_tube_ > = 1 for parallel alignment and 0 for perpendicular alignment. The thick solid line and shaded region represents mean±s.d. (F) Oil tube length (*l*_tube_) plotted as a function of time (n=18 tubes from three independent tests). The dashed line is a fit to the average tube length using an exponential equation (adjusted *R*-square = 0.96). All scale bars = 10 *μ*m.

Differential labeling of the oil and cells reveals that the tubes are not filled with water; instead, they are filled with oil (**Fig. S5A**). This result indicates that cell adhesion to the oil stabilizes the tubes against collapse, thereby preventing the deformed droplet from regaining a spherical shape. Furthermore, careful inspection of the confocal images of the tubes reveals that the bacteria are well aligned to the tube axis, which becomes clear in the orientation director field of the cells on the tube (**Fig. 2D**). We characterize cell alignment along the tube by calculating the local average scalar product between the director field and the tube axis, finding values close to 1 along the tube; this indicates that the cells are highly aligned with the tube axis (**Fig. 2E, Fig. S5B and C**, and **SI - Methods**).

Due to this parallel alignment of cells on the tubes, we hypothesize that the rate of increase of the tube length is proportional to the number of cells on the tube, which should increase exponentially if all cells were to divide. We measure tube elongation on different droplets, finding that it increases rapidly and in a manner consistent with exponential elongation, supporting this hypothesis (**Fig. 2F, Fig. S5D**, and **SI-Methods**). Furthermore, from the fit to our data, we extract a tube length-doubling time of ~3.4 h, which is 2-fold larger than the cell division time (*t*_div_) of 1.65 h (**Fig. S1D**). This difference likely arises from the imperfect alignment of cells along the tubes and from the expulsion of cells from the tube interface during elongation, which is visible around the tubes (**Fig. 2C**).

### Biofilm phenotypes are associated with a decrease of interfacial tension

The large difference in biofilm morphology between the two phenotypes suggests differences in their interfacial properties. *A. borkumensis* are known to secrete biosurfactants both on the cell surface and into the environment, which are thought to aid in the assimilation of oil (*9, 35*). These biosurfactants can lower the oil-water interfacial tension (*γ*), making it easier to deform the interface, and form conditioning films at interfaces, which can facilitate surface colonization. To measure differences in interfacial properties based on biofilm phenotype, we fractionate SB and DB cultures into three components: cells, conditioned media, and conditioned C16 to independently measure *γ* for each (**Fig. S6A**). We find that *γ* for each of the respective fractions are depressed relative to control values; however, the DB-conditioned oil decreases the most. The *γ* for DB-conditioned oil decreases from 30 to 8 mN/m and is about half the value for SBs (**Fig. 3A, Fig. S6B-D** and **Table S1**). Surprisingly, when we microfluidically “re-sample” the biofilm phenotype using the DB-conditioned C16 instead of fresh C16, we find no change in observed phenotype (**Fig. S6E**). Thus, despite the ~4 fold lower *γ* of DB-conditioned oil, the SB cells are unable to deform the oil-water interface, indicating that a lower *γ* is not sufficient to produce the DB phenotype.

**Fig. 3:**
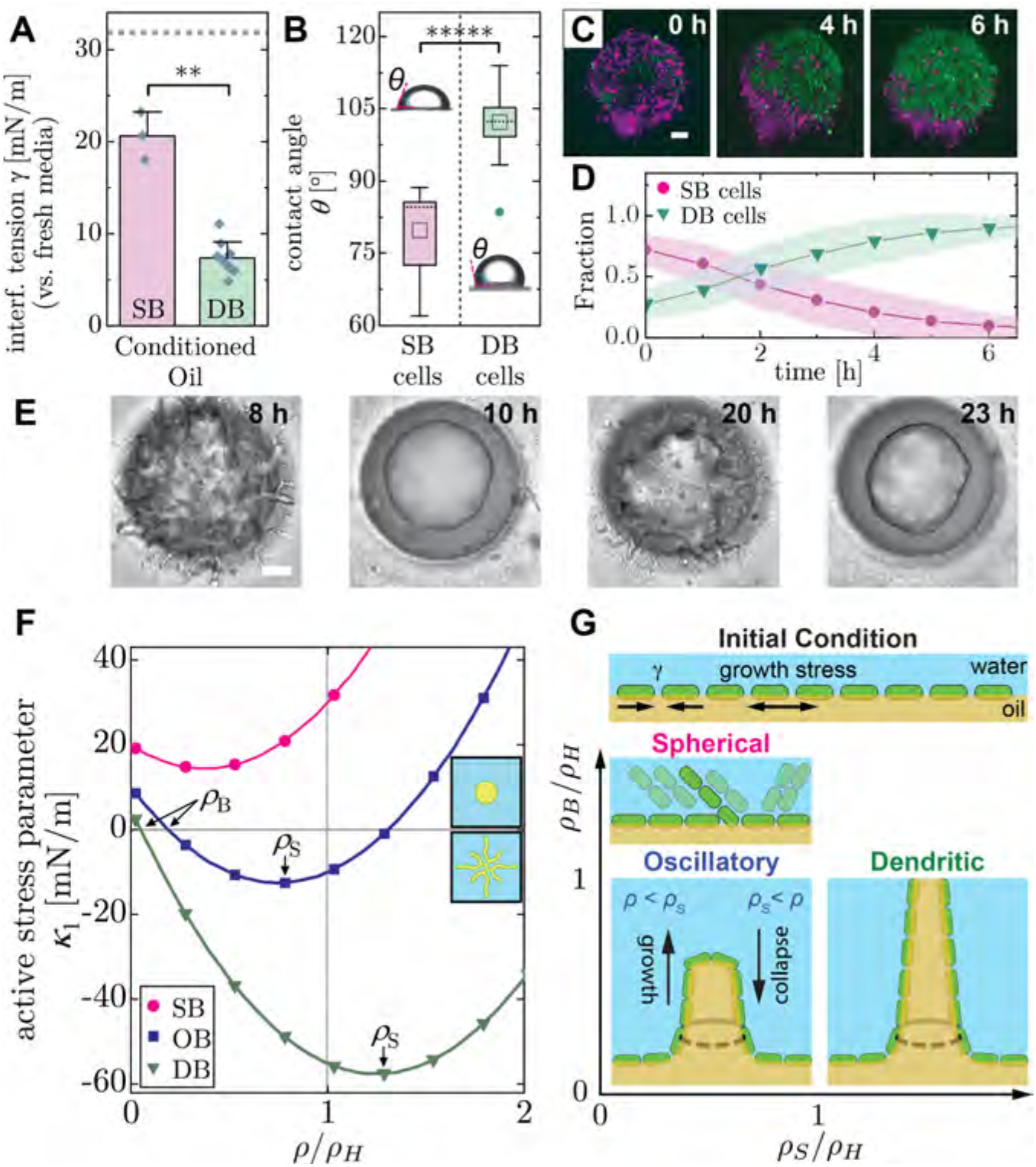
Theoretical model of tubulation. (A) Interfacial tension (*γ*) between biofilm-conditioned oil and fresh media measured using pendant drop tensiometry for spherical (SB) and dendritic biofilms (DB), respectively (from n_SB_=3 and n_DB_=10 independent tests). The dashed line indicates *γ* between fresh oil and fresh medium (n_control_=5). We used Welch’s *t*-test to compare the means (** denotes *p*<0.01; *p*=0.0062). (B) Three-phase contact angles (***θ***) between a water drop deposited on a bacterial lawn of SB and DB phenotypes, submerged in oil. The (□) is the mean, the horizontal dashed lines are the medians, and (●) represents an outlier (from n_SB_=10 and n_DB_=13 independent tests). We used Welch’s *t*-test to compare the means (***** denotes *p*<0.00001; *p*=1.2⨉10^−6^). (inset) Images of the water drops wetting bacterial lawns that are submerged in oil. (C) Time-lapse sequence showing the competition between SB cells (magenta) and DB cells (green) for interfacial area on a trapped droplet. Droplets are generated in a suspension containing SB and DB cells at a ratio of 3:1 (see **Supplementary Information** for details). DB cells displace a monolayer of SB cells over the course of ~6 h. (D) Measurement of the fractional coverage of SB and DB as a function of time from confocal images. The solid line and filled regions represent mean ± s.d. (from n=10 representative drops). (E) Oscillatory biofilm (OB) behavior demonstrated by a culture sampled for an intermediate duration between the SB and DBs. (F) Model of the tension (*k*_1_ in the phase-field model) experienced by the tip of a tube as a function of normalized interfacial cell density (*ρ*/*ρ*_H_) (see **Supplementary Information** for details). We normalize cell density by the homeostatic interfacial cell density (*ρ*_H_). Positive (negative) values of *k*_1_ indicate tube retraction (expansion). For the SB phenotype, tension is positive for all densities, thus tubes are unable to form. For the intermediate OB phenotype, the homeostatic density is located in a regime where the tension is increasing as a function of biofilm density. As cell density approaches *ρ*_H_, the increasing tension further compacts the cells, which further increases cell density, driving positive feedback that eventually triggers collapse of tube structures. The slope at the y-intercept is *k*_1_ = 3, 6, and 10 for the SB, OB, and DB models, respectively. The *ρ*_B_ is the critical buckling density, where *k*_1_<0. For SBs, *ρ*_B_=_∞_. The *ρ*_S_ is the optimal cell density, where *k*_1_ reaches a minimum, and beyond which the effect of spontaneous curvature is reduced. (insets) Phase-field simulation showing a circular droplet (*k*_1_>0) and a tubulated droplet (*k*_1_<0). (G) Schematic phase diagram of the biofilm phenotypes in terms of normalized densities: *ρ*_S_/*ρ*_H_ and *ρ*_B_/*ρ*_H_. When *ρ*_B_/*ρ*_H_ > 1, SBs form because the interfacial cell density can never become sufficiently large to induce buckling. For *ρ*_B_/*ρ*_H_ < 1, cell division drives an increase in cell density beyond *ρ*_B_. When *ρ*_S_/*ρ*_H_ < 1, oscillations between the spherical and dendritic phenotypes can occur (OB). When, *ρ*_S_/*ρ*_H_ > 1, stable dendrites (DB) occur.

Interfacial behavior is also affected by cell hydrophobicity, which, together with *γ* controls the extent to which cells are wetted by the oil (**Fig. S7A**). Cell hydrophobicity is thought to increase the longer cells consume oil, however the relationship with phenotype is unclear (*20, 36*). Although the bacteria appear to lay flat on the surface at *t*_0_, the microscopic contact angle is difficult to estimate from confocal images (**Fig. S7B)**. Thus, to estimate hydrophobicity for the cells of both phenotypes, we measure the 3-phase contact angle (*θ*) between water that we deposit on a bacterial lawn submerged in oil (*37*) (**Fig. S7C-E** and **SI-Methods**); *θ*=90° is the neutral wetting condition and larger *θ’*s indicate greater hydrophobicity. SB cells, which are isolated after 1 d of culture, have a *θ*≈80°, whereas DB cells, which are isolated after 5 d, have a *θ*≈100° (**Fig. 3B** and **SI-Methods**). This indicates that the midplanes of SB and DB cells are ±10% above and below the interface, respectively (**Fig. S7A**).

The higher hydrophobicity of DB cells indicates that they have a larger interfacial adhesion strength than SB cells. To directly compare the respective adhesion strengths of SB and DB, we force them to compete for interfacial area on oil microdroplets, noting that both phenotypes have similar division times (**Fig. S1D**). We generate cell-laden droplets using mCherry-expressing SB cells and GFP-expressing DB cells in a 3:1 ratio, and record fluorescence intensity as the biofilm develops. Since the same oil substrate is used, we ensure that both phenotypes experience the same *γ*. We find that although the DB cells are initially in the minority, they gradually displace the established SB, to become dominant in ~5 h (**Fig. 3C,D**). This result confirms our expectation that DB cells, which are more hydrophobic do indeed have a larger adhesion energy to the interface than SB cells.

Rod-shaped gammaproteobacteria like *Pseudomonas aeruginosa* have been shown to colonize and remodel oil-water interfaces with their biofilms, similar to what we observe in the early stages (<3 h) of DB formation (*17, 18*). However, those biofilms lack the large-scale deformations that we observe in later stages of DB formation (**Fig. 1C**, >5 h). From our measurement of the oil-water interfacial tension, the biofilm compression modulus is estimated to be ~200 Pa (*38*), which is much smaller than the ~1 MPa extensional growth pressure of *P. aeruginosa* (*39*). Assuming that *A. borkumensis* have a similar growth pressure, cell division supplies enough stress to easily deform the interface. Thus, DBs generate the biofilm phenotypes we observe only if a sufficient number of cells remain adhered to the interface to drive the tubulation process. Conversely, the lack of deformations in the SB phenotype is the consequence of cell detachment from the interface followed by biofilm formation around the droplet.

### Membrane theory predicts the transition from SB to DB phenotype

Based on these observations, we develop a coarse-grain membrane model to describe the interfacial dynamics of the growing interfacial biofilm. The model explains the transition between SB and DB phenotypes in terms of a competition between the interfacial tension and the spontaneous curvature of the biofilm, which is its intrinsic tendency to bend in a preferred direction. Interfacial tension resists expansion of the surface by the biofilm, while the spontaneous curvature of the biofilm governs the shape of the expanding surface. When the energetic cost of increasing surface area is lower than the cost of bending, tubes are generated, which expand exponentially according to the following equation:

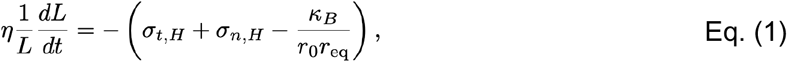

where *L* is the tube length, *η* is the viscosity of the biofilm layer, *σ*_*t,H*_ and *σ*_*n,H*_ are the respective tensions along the circumferential and normal directions of the tube, *κ*_B_ is the bending rigidity of the membrane in the circumferential direction of the tube, 1/*r*_0_ is the spontaneous curvature of the biofilm, and *r*_eq_ is the tube radius (**SI – Membrane Model** and **Fig. S14**).

**Eq. 1** encompasses the existence of active extensile nematic stresses driven by bacterial growth, with an effective interfacial tension *γ*_eff_(*ρ*) = ½(*σ*_*t,H*_ *+ σ*_*n,H*_) that depends on the interfacial cell density *ρ* (**SI – Membrane Model**). This implies that tubulation occurs at a critical buckling density (*ρ*_B_), consistent with our observation that tubes form after confluency (**Fig. 1C**). In our model, bacteria populate a circular oil-water interface with a density that increases logistically towards a homeostatic density (*ρ*_H_), where division and loss are balanced. Depending on the ratio *ρ*_B_/*ρ*_H_, the phenotype changes: when *ρ*_B_/*ρ*_H_ > 1, tubes are unable to form, resulting in the SB phenotype; when *ρ*_B_/*ρ*_H_ < 1, the interface buckles and stable tubes form, producing the DB phenotype. Our model implies that physically lowering *ρ* by removing cells of a DB below the critical value should cause a deformed droplet to recover its unperturbed spherical shape. To test this hypothesis, we expose DBs to a flowing surfactant mixture to disrupt the biofilm (**SI - Methods**). Following a ~4 h lag, the biofilm abruptly washes away and the deformed droplets become spherical; this recovery is the consequence of positive interfacial tension in the absence of bacteria (**Fig. S8A,B** and **Movie S3**).

In addition to the SB and DB phenotypes described so far, we observe an intermediate phenotype when we isolate cells at an intermediate culture time (**Fig. S1A**). These biofilms present dynamic oscillatory behavior (OB), alternating between the dendritic and spherical biofilms, with a period of ~12 h (**Fig. 3E, Fig. S9**, and **Movie S4**). The relatively short (~30 min) transition from tubulated to spherical is consistent with a sudden loss of tube stability, which, based on **Eq. 1**, is consistent with a sudden increase in the effective surface tension.

To elucidate the emergence of oscillations between spherical and dendritic phenotypes, we use a phase-field approach following an established literature on simulating interfacial dynamics of multiphasic systems (*40*). In our case, the simulated field is the local fraction of oil (*φ*(x,y)) that obeys a Cahn-Hilliard equation with an interfacial term *κ*_1_(*ρ*)(*∇φ*)_2_/2; here, *κ*_1_(*ρ*) is defined by right hand side of **Eq. 1** and sets the tube elongation rate. *κ*_1_(*ρ*) is thus a proxy for the active stress along the interface (**SI - Membrane Model**). In our model, a linear relation of the form *κ*_1_(*ρ*) = *γ* - *k*_1_ *ρ*, where *k*_1_ accounts for the force exerted by the biofilm on the oil is sufficient to understand the SB-to-DB transition. The interface remains spherical until *ρ* surpasses the critical density, *ρ*_B_ = *γ* / *k*_1_; here, *κ*_1_(*ρ*) becomes negative and tubulation occurs as the interface buckles. However, the morphological oscillations emerge only if we consider a second-order expansion of *κ*_1_(*ρ*) in *ρ*, such that an ‘optimal’ cell density exists where tube elongation and the final tube length are maximal, which we denote *ρ*_S_ (**Fig. 3F**). Oscillations arise for the specific case when *ρ*_B_<*ρ*_S_<*ρ*_H_ (**SI - Phase-Field Model**); here, as cell density increases beyond the optimal value *ρ*_S_, tube elongation ceases, and contraction starts. During this contraction, the surface shrinks faster than cells can be ejected from the interface, leading to an abrupt increase in bacterial density. We find that *ρ* overshoots *ρ*_H_ and causes *κ*_1_(*ρ*) to become positive (**Fig. 3F, blue**). This dynamical overshoot ultimately induces a catastrophic collapse of the tubes and is accompanied by a large reduction in the number of cells at the interface. Consistent with this prediction, in our experiments we observe a large and persistent flow of cells away from the interface soon after tube collapse (**Movie S4**). Our biomechanical model recapitulates the transition from these three phenotypes in terms of a tubulation mechanism that depends on bacterial growth dynamics, with oscillations emerging through a dynamical phase transition mechanism (*41*) (**Fig. 3G, Movie S5**). These oscillations are driven by continuous cell division that pushes density beyond the critical values for tube growth and collapse.

### Controlled Buckling of Confined Droplets

We leverage microfluidics to position the topological defects that generate tubes by controlling the shape of the trapped drops. By using droplets whose diameters are larger than the trap height, we can generate biofilms on drops that are flattened by the trap ceiling (**Fig. 4A - Upper inset)**. These flattened circular faces allow us to position the nematic defects in the center, which concentrates growth stress and thereby generates centrally located dimples. Unlike smaller drops, which are uniformly covered with a bacterial monolayer, interfacial tension initially excludes bacteria from the flattened regions on large drops (**Fig. 4A** and **Fig. S10(A-B)**). At a critical cell density, bacterial growth pressure enables cells to invade the flat regions, which they do so isotropically (**Movie S6**). Once confluency is reached, the nematic field displays an aster-like +1 defect (or two closely spaced +1/2 defects) in the center, as expected from liquid crystal theory (*42, 43*) where flows orient the cell nematic director (**Fig. 4B** and **Fig. S10(C-D)**). As cell division continues, we observe the formation of a large dimple at the defect, which grows inward into the droplet (**Fig. 4C** and **Fig. S10E)**. By symmetry, a similar process occurs at the top of the drop (**Fig. S10B-inset**). We find that the height profile of the dimple agrees with a model of spontaneous deformation of a liquid crystalline membrane with a radial nematic order (*30, 44*), given by:

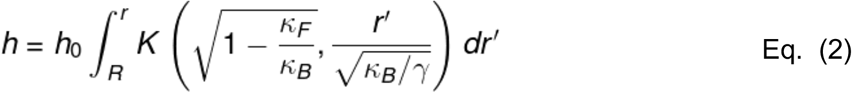

where *h* is the dimple height, *r* is the radial position, *R* is the radius of the dimple, *K*(***α***,*x*) is a modified Bessel function of order *α*, and *κ*_*F*_ is the elastic constant of the nematic director field (see **SI – Fitting Procedure**). The theoretical profiles depend on a scaling factor *h*_0_ and the two dimensionless parameters *k*_*F*_/*k*_*B*_ and *k*_*B*_/*γR*^2^; these parameters arise from the competition between the Frank free energy and the bending energy, and between the bending energy and the surface tension, respectively. Our fitting procedure yields a ratio *k*_*F*_/*k*_*B*_ ≈ 2, highlighting the prominent role of the bacterial nematic order on the dimple profile (**Fig. S12**). In agreement with experiments, the profiles in this parameter regime are qualitatively similar to pseudospheres (**Fig. 4D** and **Fig. S13**). The rounded shape at the inward tip is a consequence of the finite size of the bacteria, which have non-zero bending moduli.

**Fig. 4:**
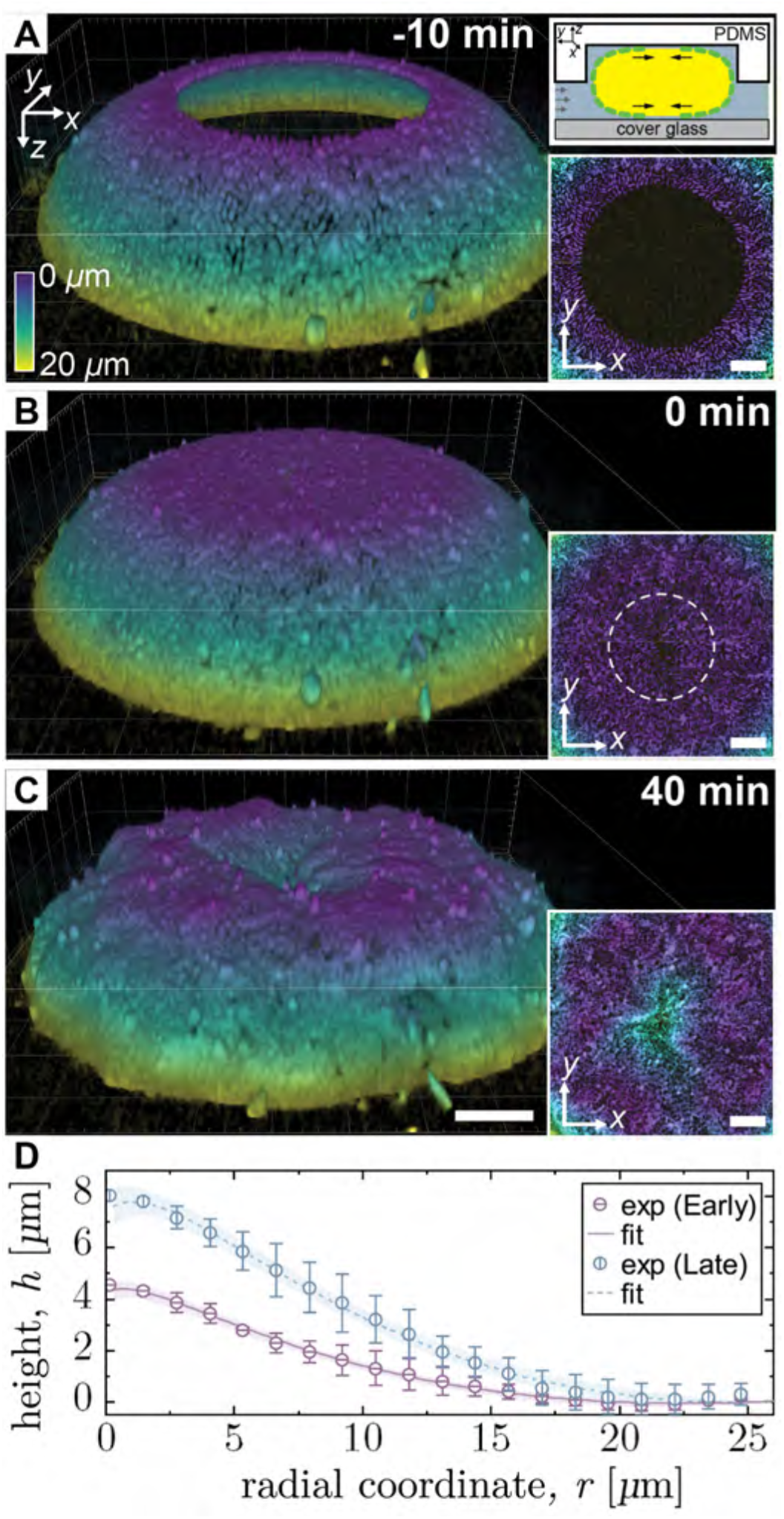
Defect mediated buckling of the surface of a confined droplet. (A-C) Time-lapse confocal image sequence showing the evolution of a dendritic biofilm on a ‘flattened’ drop, color coded by depth. (A) Prior to confluency (−10 min), interfacial tension excludes cells from the flattened regions at the top and bottom of the droplets. These regions are generated where the non-wetting droplets make contact with the glass floor and PDMS ceiling. See the (upper) inset for a droplet schematic and the (lower) inset for the top view of the droplet. (B) Confluent monolayer is formed at *t*_0_ = 0 min (see inset and **Fig. S10**). (inset) Top view of droplet with a circle enclosing the location of the nematic defect. (C) Dimple formation in the droplet at the defect. All scale bars = 10 *μ*m. (D) Evolution of the dimple height (*h*) from *xy*-line profiles measured across the droplet midplane, at 20 min (early) and 26 ± 3 min (late) post-confluency. Dimple height is shown as a function of the radial coordinate (*r*). The fits to the data are based on **Eq. (2)**. The solid error bars and filled regions represent ±s.d on the experimental data and fits, respectively (see **Supplementary Information** for details on the fitting procedure and error estimation). Early-profile data are the average of the *x*- and *y*-profiles (n=4) at *t* = 20 min, while the late-profile data are the average of the *x*- and *y*-profiles binned at 23, 26, and 29 min (n=12), respectively. To generate the dimensional values shown, we rescale the non-dimensionalized *h*-values by the dimple heights, 4.5 *μ*m and 8.0 *μ*m at early and late times, respectively, while *r* is rescaled by 26 *μ*m at all times.

## DISCUSSION

Although bacteria are known to play an integral role in the bioremediation of spilled oil, the microscopic mechanism of how hydrocarbonoclastic marine bacteria engage and break-down oil remains largely unexplored. The process of biodegradation has been studied using *in situ* microcosm tests and batch reactors, which cannot reveal the underlying physics at the oil-water interface. To circumvent the vagaries of bulk and average measurements, we leverage oil-drop microfluidics to enable simultaneous longitudinal tracking of numerous individual oil-consuming biofilms, at unprecedented spatial and temporal resolution. By sampling *A. borkumensis* cells from cultures that have been adapted to metabolizing oil for variable durations, we show that the expressed phenotype depends on the culture duration and is more complex than previously known. Cells adapted for oil consumption possess the ability for explosive growth, generating biofilms that are morphologically distinct from biofilms formed from less-adapted cells.

We explain the appearance of this optimized phenotype by linking it to measurable increases in cell hydrophobicity and interfacial adhesion, which correlate to cultivation time with oil. These optimized cells collectively manipulate the shape of their carbon source to substantially increase the number of cells that may simultaneously feed; this greatly increases the rate of oil consumption. And, although ‘higher’ organisms enact similar food-reshaping activities (*45*), to the best of our knowledge this type of cooperative behavior is unique in bacteria. Surprisingly, at the single-cell level, the optimized cells are no more efficient at consuming oil than are their unoptimized counterparts, despite their considerably more efficient biofilms. Taken together, this increase in efficiency may be one factor, among many, that leads to the observed blooms of *A. borkumensis* and other similar organisms after oil spills (*46*).

The high spatio-temporal imaging of numerous optimized biofilms afforded by our platform, further reveals the mechanism of their interfacial manipulation: as cell density increases past confluency, the biofilm buckles at aster-like defects in the nematic field, allowing the cells to escape the confines of the droplet surface. Importantly, rather than peeling away from the oil, the biofilm protrusions entrain oil as they elongate into tubes. Recent developments on defect-driven morphogenesis have highlighted the importance of defects in the nematic order (*26, 28, 30, 32, 33*). We show indeed that *A. borkumensis* biofilms utilize nematic defects to enact topological deformations of the interface that results in exponentially elongating tubes, generating new surface area where more cells may feed. Thus far, models of oil biodegradation have treated droplets as shrinking spheres. Here, we develop an active matter theory to capture the dynamics of tube formation based on the most recent development of active nematics on curved geometries (*47*). Our model recapitulates each phenotype and can explain the existence of the striking oscillations in the biofilm morphology. Since we obtain OB cells at intermediate adaptation times, these cells may present an intermediate hydrophobicity. It will be interesting to explore whether a coupling between growth and competition for resources between un- and partially-adapted cells drives the oscillatory behavior (*48*).

In one set of experiments, we show that addition of surfactants that mimic oil-dispersant formulations to the culture medium leads to rapid biofilm detachment from the oil; this uncoupling abruptly halts oil consumption. Other than screening for enrichment of specific microorganisms in microcosm tests (*13*), to our knowledge, there is no experimental way to directly assess how medium composition affects oil-degradation by OHCB, despite the practical importance of this question. Our result clearly shows that surfactants can significantly affect biofilm formation; however, in the environment, many factors can possibly affect the final community composition, such as nutrient concentration, surfactant dilution, the specific mechanism of interfacial adhesion by different microbial species, and surfactant tolerance. Thus, how surfactants shape OHCB biofilm development remains an open question.

Application of this method to analyze other OHCB species could begin to provide information that complements metagenomic screens and could serve as a starting point to investigate how OHCB behave under different conditions. Our microfluidic platform is sufficiently general to accommodate studies of the population dynamics of both mono- and mixed-cultures of oil-consuming microorganisms subjected to different oil substrates, culture conditions, and different oil-dispersants. Results from such tests could provide insight into improving current remediation procedures in the field ranging from guidance on the suitability of a specific OHCB to a particular environment, to the potential of pre-adapting organisms of interest to local conditions to accelerate bioremediation.

## CONTRIBUTIONS

MP, NO, KS performed the experiments; MP, NO, NN, JF, ASU analyzed and discussed the data; CBM, JP, JFR, SZL designed the tubulation theory; SZL carried out simulations; CBM performed the dimple height analysis; MP, CBM, NO, JFR, JF, ASU wrote the manuscript; JF and ASU conceived of the project.

## ACKNOWLEDGMENTS

We thank Y. Yamashita (UT) for use of the contact angle/pendant drop systems, F. Pincet (Laboratoire de Physique, ENS) for assistance with micropipette experiments, D. Quéré (ESPCI) for the fruitful discussion on the O/W/cell contact angle measurements, and F. Brochard-Wyart (Institut Curie) for discussions on tube formation.

## FUNDING

The research leading to these results was supported by the JSPS–ANR SAKURA Grant (JF, ASU, NO), JSPS BRIDGE Fellowship (JF), JST-ERATO (NN) (JPMJER1502), JSPS Kakenhi B (ASU) (21H01720), France 2030 (ANR-16-CONV-0001) (JFR), Excellence Initiative of Aix-Marseille University - A*MIDEX (JFR) and ANR-20-CE30-0023 (JFR).

## COMPETING INTERESTS

The authors declare that they have no competing interests.

## DATA AND MATERIALS AVAILABILITY

All materials are available upon request.

## SUPPLEMENTARY MATERIALS

Supplementary Information:

- Materials And Methods,
- Analytical Model of Oil Consumption
- Extended Tubulation Model
- Fitting Procedure for the Dimple Profiles
- Supplementary Figures: Fig S1-S17
- Supplementary Tables: Table S1-S2
- Supplementary Movies: Movie S1-S6
- Supplementary References (*1-29*)

## Notes

### Competing Interest Statement

The authors have declared no competing interest.

### Summary of Updates

Final preprint before submission

