## Supplementary Information for "*Alcanivorax borkumensis* Biofilms Enhance Oil Degradation By Interfacial Tubulation"

**S9 Supplementary Figures** 24

**S10 Supplementary Tables** 41

**S11 Supplementary Movies** 42

### S1 Bacterial methods

#### S1.1 Solid and liquid culture methods

We purchased *Alcanivorax borkumensis* SK2 bacterial strain from ATCC (ATCC® 700651TM). Cells are cultured by streaking from a freeze stock onto Difco Marine Agar Plates (BD Difco; Marine Agar 2216) supplemented with 10 g/L (90 $\mu$M) sodium pyruvate (Sigma-Aldrich; P2256; CAS#113-24-6), a known water-soluble carbon source [1]. After incubating the solid cultures at 30°C for 2 days, we select ~5 isolated bacterial colonies and inoculate in 4 mL of Marine Broth (MB) (BD Difco; Marine Broth 2216) supplemented with 10 g/L of sodium pyruvate, which we refer to as MB+pyr. We incubate the liquid culture in a linear axial shaker (Taitec, Japan) at 190 rpm at 30°C for 2 days and then wash the cells 2x in ONR7a (DSMZ medium #950, DSMZ, Germany), a chemically defined artificial seawater medium. We measure the optical density of liquid cultures at a wavelength of 600 nm ($OD_{600}$) using a Nanodrop spectrophotometer (Thermo-Fisher scientific, Nanodrop 2000/2000c UV-Vis).

After washing the cells, we resuspend them in ONR7a at $OD_{600}$=0.01, and supplement with 100 $\mu$L of hexadecane (C16) (Sigma-Aldrich, 296317, CAS #544-76-3); this medium is referred to as ONR7a+C16. This volume of C16 has a concentration of ~85 $\mu$M in 4 mL, which is close to the concentration of sodium pyruvate in MB+pyr. These steps are summarized in the culturing box in **Fig. S1A** and growth curves for *A. borkumensis* SK2 grown in ONR7a+pyr and ONR7a+C16, respectively, are shown in **Fig. S1B**.

To measure the $OD_{600}$ for cultures containing oil, we first allow the emulsion to rise by storing the test tubes statically for 5 min. Next, using a 'long' pipette tip, we aspirate 100 $\mu$L of the cell suspension from the bottom of the test tube and transfer it to a cuvette for the OD measurement.

#### S1.2 Phenotype Identification

We adapt the bacteria to growing on C16 by incubating this liquid culture for a period of 1-5 days, under the shaking conditions described earlier. Over this time, distinct phenotypes emerge, which we identify based on the drop deformation dynamics in the microfluidic devices (**Fig. S2**). The spherical biofilm (SB) phenotype appears after 1 d of shaking culture (**Fig. S3B**), while the dendritic biofilm (DB) phenotype appears after 5 days (**Fig. S3C**). At an intermediate period of time of 3-4 d, we find an intermediate state that possesses characteristics of both the SB and DB phenotypes, such as thin biofilms that cause only small-scale deformations, relatively short oil tubes, and an oil degradation rate similar to SBs. At times, the presented phenotype may oscillate between the two phenotypes. For this reason, we designate this mixed state the oscillatory biofilm (OB) phenotype (**Fig. S9**). We are unable to quantify the relative ratios of SB or DB cells in this phenotype.

We find that OBs and DBs develop from ~25% of the cultures. The adaptation, microfluidic sampling, and phenotype isolation processes are summarized boxes 2-5 in **Fig. S1A**. To ensure sufficient sampling we typically use about 10 test tubes to select the OB or DBs. In addition to differences in oil degradation dynamics, the respective test tube cultures also show differences that are evident by inspection if they are cultured for 5 days (**Fig. S1C**). The SB phenotype is characterized by a relatively lower $OD_{600}$ liquid culture with a clearly visible biofilm attached to the tube wall near the bottom. The OB and DB phenotypes are characterized by higher $OD_{600}$ cultures that lack the visible biofilms and with smaller emulsion drops than the SB tubes. The size of the emulsion drops in DBs tubes, however, are significantly smaller than OB emulsions, as shown in **Fig. S1C**.

#### S1.3 Fluorescent Strain Construction

We construct fluorescent reporter strains by transforming *A. borkumensis* SK2 with plasmids containing either $P_{gap}$-gfp or $P_{gap}$-mCherry, which constitutively express GFP or mCherry, respectively [2]. When culturing fluorescent trains, we include 125 $\mu$M gentamicin in the media to prevent the loss of the fluorescent plasmid.

### S2 Microfluidics

#### S2.1 Device fabrication and channel coating

We fabricate two-layer microfluidic devices using two-step soft-lithography techniques [3]. Briefly, we spin coat the first layer of KMPR photoresist (Microchem, MA, USA) onto a silicon wafer and then expose to $i$-line UV radiation through a photomask; this forms the channel layer. We then spin-coat a second layer of photoresist on top of the first layer, align a second photomask to the alignment marks in layer 1, and then expose a second time; this forms the traps. Development of the unpolymerized KMPR reveals a two-layer master mold. We then pour well-mixed PDMS containing curing agent (Dow Corning, Sylgard 184) in a 10:1 ratio onto the silicon master, degas, and then bake overnight at 75°C. We carefully peel the hardened PDMS slab from the master mold, punch holes for inlet and outlet channels, and remove debris using pressurized N2 gas. We treat the PDMS slab and cover glass using oxygen plasma (CUTE-MPR, FemtoScience, Korea) for 30 s at 100 W and then bring the two into contact with each other, which then bond permanently. A schematic of the filled device is shown in **Fig. S2A**.

Since PDMS readily absorbs organic solvents, we coat our devices with polyvinyl alcohol (PVA) (Sigma-Aldrich; 363170; CAS #9002-89-5) to create a barrier that prevents absorption of C16 [4] . To generate this coating, immediately after plasma bonding we degas the devices under vacuum while submerged in a 2 w/w% PVA dissolved in deionized filtered water (milliQ) solution for 2 h. We then flush the device with pressurized N2 gas to remove the liquid and bake for 15 min at 110°C. We repeat this procedure 4x and then use the devices directly or store them submerged in a 0.2 w/w% PVA solution. This coating fully suppresses any measurable absorption of C16 by PDMS for up to 3 weeks. The radii of 15 trapped drops are measured as a function of time and shown in **Fig. S2B**.

#### S2.2 Device operation

To facilitate long-term biofilm imaging on oil drops, we incubate the cell-laden oil drops in a two-layer microfluidic device that utilizes surface tension to trap them in vertically-oriented cylindrical pockets [5]. We load the device with cell-laden drops that are injected through the drop inlet (see **Fig. S2A**). Drops whose diameters are larger than the height of the main channel are flattened into disks, while smaller droplets remain spherical. When the flattened droplets encounter a vertical pocket, they are able to to reduce their surface area-to-volume ratio by becoming more spherical; this traps the droplets, as shown in **Fig. S1A**. After trapping, the flow through the droplet inlet is closed using a three-way valve, and we then infuse media at a flow rate of 30 $\mu$L/h through the media inlet using a syringe pump.

We generate bacteria-laden droplets for microfluidic sampling by harvesting cells grown on ONR7a+C16, washing twice in ONR7a to remove the oil, and then resuspending at a final $OD_{600}$ of 0.75-1.0 containing 0.1 wt% Tween 20 (Sigma-Aldrich; P9416; CAS# 9005-64-5). To generate this $OD_{600}$, we typically culture 4 test tubes. We then pipette 100 $\mu$L of fresh C16 on top of the washed cell culture and vortex for 100 s at 3000 rpm to generate an emulsion of cell-laden drops with diameters in the range of 10-150 $\mu$m. We allow the suspension to rest for 5 min before loading into a syringe to infusion into the device (see **Step 3** in **Fig. S1A**).

Prior to introducing the droplets, we prime the device by filling with ONR7a, taking care to remove all bubbles. We then gently infuse the emulsion into the drop inlet port with a syringe pump (kdScientific LEGATO 200-788200). Some fraction of the drops become trapped while the rest are washed away. After filling the traps, we close a three-way valve that is connected to the droplet inlet to prevent backflow. This method enables us to trap droplets with $\sim$20-50 attached cells in nearly all the traps. We then wash the Tween 20 from the device by pumping fresh ONR7a through the media inlet for 2 h at a flow rate, Q, of 50 $\mu$L/min [6].

Immediately prior to image acquisition we set Q = 0.5 $\mu$L/min; this flow rate replenishes the volume of media in the device once per minute. A confluent bacterial monolayer typically develops over the course 12-14 h depending on the initial cell density. A time lapse sequence capturing this process is shown in **Fig. S3A**. When combined with bacterial division time and drop radius, we estimate the initial number of attached cells to be ~20-50 cells. We assign the time of monolayer formation as $t_0$ for all experiments.

##### S2.3 Cell division time

For the cells grown using pyruvate, which is directly soluble in media, we use a high-aspect ratio microfluidic device to facilitate the imaging of individual cells [7]. We determine the division time, $t_{div}$, by recording the interval between division events from individual cells for different generations and across different lineages. We determine $t_{div}$ of the isolated phenotypes by recording the interval between division events of cells attached to trapped oil droplets. The division times for cells cultured in pyruvate is 1.74 $\pm$ 0.26 h, SB cells is 1.76 $\pm$ 0.21 h , and DB cells is 1.64 $\pm$ 0.23 h, as shown in **Fig. S1D**.

##### S2.4 Fluorescent labeling of oil

To label the oil drops, we mix the hydrophobic dye FM 4-64 (Thermofisher, CAS# : 162112-35-8) into the media at a final concentration of 2.5 $\mu$M and incubate at the start of the experiment. This dye has the added benefit of labeling the cell membrane.

#### S3 Interfacial Properties

##### S3.1 Pendant drop tensiometry

We determine the interfacial tension, $\gamma$, of the oil/aqueous interface under the influence of: 1) harvested cells; 2) conditioned culture media; or 3) conditioned (partially consumed) oil, respectively using the pendant drop method (KYOWA DM-305, Saitama, Japan). The rationale to test these phases is that *A. borkumensis* secrete biosurfactants that that could accumulate in the culture media or in the partially consumed C16, affecting $\gamma$ [8, 9]. In addition, *A. borkumensis* are known to secrete amphiphilic molecules, some of which remain directly on the cell surface [8, 10, 1]. Since autolysis is also thought to be important for *A. borkumensis* biofilm formation [11], when we separate the oil phase from liquid culture, membrane-bound amphiphilic molecules may remain directly on the oil, which could lower $\gamma$.

For these tests, we first prepare liquid culture of the desired phenotype and harvest the different phases. To separate the different phases (cells, conditioned media, and conditioned oil) from each other we centrifuge the culture at 10°C for 30 min, which simultaneously sediments the cells and freezes the C16 at the top of the media. We gently transfer the frozen C16, the supernatant, and the cells to new tubes and then further purify each respective phase. First, we wash the harvested cells 3x in fresh ONR7a ($10,000g$ for 30 min at 25°C) and set the final $OD_{600}$ $\approx$0.5 in 10 mL. Next, we sterilize the conditioned supernatant by filtering it through a 0.22 $\mu$m pore filter (Merck Millipore). Finally, we remove cells and cellular debris that may be attached to the conditioned oil by centrifuging ($20,000g$ at 25°C for 30 min) and transferring to new tubes three times. The separation process is summarized in **Fig. S6A**.

We then measure $\gamma$ between: 1) the harvested cells suspended in fresh media with fresh C16; 2) the cell-free conditioned media with fresh C16; and 3) the conditioned C16 with fresh media using pendant drop tensiometry. Here, we suspend a 5 $\mu$L oil pendant drop vertically in a glass cuvette (3x3x2 cm3) that has been filled with the aqueous phase, using a U-shaped needle attached to a 5 mL syringe and record images at regular intervals. Software from the instrument outputs $\gamma$. We measure $\gamma$ = 33$\pm$5 mN/m for fresh media and fresh C16. We find that for the SB phenotype, each of the fractionated phases gives $\gamma \approx$ 20 mN/m (see **Fig. S6B-D**). For the DB phenotype, the corresponding $\gamma$-values are larger than those of the SB phenotype except for the conditioned oil, where we find that $\gamma$ decreases to ~8 mN/m (**Fig. 3A** and **Fig. S6B-D**).

#### S3.2 3-phase Contact Angle Measurement

We measure the 3-phase contact angle, $\theta$, between water, cells, and oil by depositing water droplets on top of a bacterial lawn that we submerge in C16 [12, 13, 14]. To form the lawn, we first culture the bacteria under appropriate conditions to generate the desired phenotype described in the Bacterial Culture Methods section. To ensure that we have a sufficient amount of cells, we culture 10 test tubes containing 4 mL apiece with an initial $OD_{600} \sim 0.05$. We first separate bulk oil from the cultures by centrifuging (30 min at 10,000 g) and resuspending in milliQ deionized water. We then harvest the cells by washing 3x in milli-Q water (10 min at 16,000 g) and resuspending in 10 mL, setting the $OD_{600} \approx 1$. We remove any bacterial aggregates in this suspension by filtering with a 5 $\mu$m pore Durapore membrane filter (Merck Millipore Ltd.).

To form the bacterial lawn, we gently pass the cell suspension through a hydrophilic PVDF 0.45 $\mu$m pore Durapore membrane filter (Merck Millipore Ltd.) until it becomes clogged. We then carefully dry the bacteria-clogged filter horizontally for 24 h at 30°C. For control experiments, we pass 10 mL of milli-Q water through the membrane and dry it under the same conditions.

After drying, we place the filter in an empty glass cuvette (5x5x5 $cm^3$) and carefully fill it with 15 mL of C16. We then deposit a 12 $\mu$L drop of milli-Q water on the submerged bacterial lawn, and record $\theta$ using a contact angle measurement system (KYOWA DM-305, Saitama, Japan). After an initial spreading of the drop, $\theta$ plateaus over the course of a few minutes as the drop reaches equilibrium. The control experiment on the native Durapore membrane gives $=133 \pm 5°$, while $\theta$ is $80 \pm 9°$ and $102 \pm 8°$ for SB and DB cells, respectively, as shown in **Fig. 3B**. Images showing the deposited water droplet on the bacterial lawn are shown in **Fig. S7C-E**.

#### S3.3 Microfluidic Sampling of Conditioned Oil

We test the conditioned oil obtained using the fractionation process described in the Pendant drop tensiometry section to determine if the lowered $\gamma$ affects the observed biofilm phenotype. For these tests, we exclusively use oil conditioned by DBs.

We first prepare liquid culture of the desired phenotype and harvest the cells as described in the Culture section. Next, following a slightly modified protocol from that described in the Microfluidic device operation section, we generate cell-laden drops using the conditioned oil instead of fresh oil. We then sample using microfluidics, finding that inoculated SB cells exhaustively form SBs, whereas inoculated DB cells exhaustively form DBs (see **Fig. S6E**).

#### S3.4 Interface Competition Test between SB and DB cells

Following the method described in earlier, we generate SB and DB cells that constitutively express mCherry and GFP, respectively. To prevent the loss of the plasmid during replication, 125 $\mu$M gentamycin is added to the culture media. We synchronize the culture times so that both SB and DB cells are formed on the same day. Then, we wash the respective cultures once and prepare a mixed suspension with a 3:1 ratio of SB to DB cells at an $OD_{600} \approx 1$ in 5 mL. Next, we generate bacteria-laden oil droplets with the mixture as described earlier, load the drops into the microfluidic device, and begin culturing. We begin two-channel confocal image acquisition at $\sim$12 h, when the bacteria have nearly formed a monolayer. To prevent blurring of the images due to movement of droplets, we select droplets with diameters that are approximately equal to the total height of the traps. These drops are free from movement. From the confocal images, we calculate the biovolume of the different phenotypes using Imaris 9.8 (Oxford instruments).

##### S3.5 Mock Corexit formulation

Although we were unable to obtain commercial samples of Corexit 9500 (Nalco Holding Company), we prepared a surfactant mixture similar to Corexit 9500 (Place et al., 2016) by mixing surfactants bis-(2-ethylhexyl) sulfosuccinate (also known as DOSS) (Sigma-Aldrich; 323586; CAS#577-11-7) (18% w/w), Span 85 (Sigma-Aldrich; S7135; CAS#26266-58-0) (4.4% w/w), Tween 80 (Sigma-Aldrich;P1754; CAS#9006-65-6) (18% w/w), and Tween 85 (Sigma-Aldrich; P4634; CAS#9005-70-3) (4.6% w/w) following previously published estimates derived from analytical liquid chromatography [15]. We excluded an enantiomeric mixture of $\alpha$- and $\beta$-ethyl hexylsulfosuccinate ($\alpha$-/$\beta$-EHSS, 0.28% w/w) from our mock Corexit since it is unclear whether it is a component of Corexit or simply an impurity.

##### S3.6 Exposing DB biofilms to the Corexit solution

We determine the critical micelle concentration (CMC) of our mixture by measuring the surface tension of the air/liquid interface of ONR7a media containing different concentrations of the dispersants using a surface tensiometer (Kyowa DY-300, Saitama Japan). We find that the CMC = 0.001 wt% (**Fig. S8C**).

To investigate the effect of the surfactant mixture on droplet biofilms, we expose 6-8 h old DBs growing on trapped droplets to different concentrations of the mock Corexit mixture ranging from 25-100$\times$CMC. Interfacial tensions of the mock Corexit are shown in **Fig. S8D**. To precisely detect the entry time into the droplet chamber, we added 3 $\mu$M FITC (Sigma-Aldrich, 34321-M, CAS No. 3326-32-7) to the culture media and acquired bright-field and fluorescent images at the outlet of the device (see **Fig. S2A** for port location). This method allows us to use media fluorescence as a proxy for Corexit concentration, as shown in **Fig. S8E**. Approximately $\sim$100 min is required for the steady state concentration in the device to be reached after infusion.

#### S4 Imaging and Analysis

##### S4.1 Bright-field and confocal image acquisition

We track spherical and dendritic biofilm development on C16 droplets by acquiring either bright-field (Zeiss Axio observer) or confocal (Zeiss LSM780 or Olympus SPINSR10) time-lapse sequences over the course of 1-4 days. Both the LSM780 and SPINSR10 have enhanced resolution modes, called Airy scan or confocal super-resolution , respectively. We typically use 63x or 100x oil immersion lenses with numerical apertures (NA) of 1.40 or 1.46. Confocal images are rendered using Imaris 9.8 (Oxford instruments). Unless explicitly stated, microscopy images in the main text are maximum intensity projections of the acquired stacks.

During confocal imaging, we use strains that constitutively express fluorescent proteins EGFP and mCherry or we label the bacteria with fluorescent membrane dyes such as FM 4-64 or FM 1-43 (Thermo-Fisher Scientific; F34653) by adding it to culture medium flowing in the device. To visualize WT cells at the oil-water interface, we add fluorescein isothiocyanate (FITC) (Thermo-Fisher Scientific; F1907; CAS#3326-32-7) at a concentration of 0.5-1 $\mu$M to the medium. FITC readily labels the cells but has low affinity for C16.

##### S4.2 Drop radius, surface area, and volume measurements

For SBs, since for the shape of droplets remains approximately spherical until $\sim$72 h, we measure the droplet radius ($R$) at its equator and use it to calculate surface area $S(t) = 4\pi R(t)^2$ and droplet volume $V(t) = \frac{4}{3}\pi R(t)^3$. For DBs since the droplet deforms soon after monolayer formation, the drop radius is ill defined. To estimate surface area, using Imaris, we draw closed contours along the fluorescent interface in each plane of confocal images and combine these using the 'surface' function to create a surface. We measure surface area from this object.

We calculate $S$ and $V$ as follow: $S = \sum_0^m P_i \Delta z$ and $V = \sum_0^m A_i \Delta z$, where $m$ is the number of $z$-plane slices that span the droplet, $P_i$ is the perimeter of $i$-th contour line, $\Delta z$ is the height resolution in the $z$ direction, and $A_i$ is the $i$-th contour line enclosing an area. $\Delta z$ is typically $\sim$ 1-2 $\mu$m.

#### S4.3 Generating a surface from bright-field image sequences

We estimate the shape from a set of bright-field images using algorithms written in MATLAB [16, 17]. These algorithms determine the focus for each pixel in a local window in an image sequence with different focus, which is then used to reconstruct the shape. For the focus measurement, we use a gradient-based operator that calculates focus from the first derivatives of the image with the assumption that focused images present sharper edges than those of out-of-focus images. The shape reconstruction technique is based on the Gaussian model of defocus [16]. In this technique, pixels with the highest focus values are identified in the image, and then a depth map is constructed by interpolating a Gaussian function around this pixel. A local focus window size of 1.5-2 $\mu$m generated realistic drop shapes.

#### S4.4 Nematic order and defect detection

Topological defects are regions in the nematic director field where the local nematic order is not defined. Topological defects are characterized by a charge, which is calculated by the number of times the director field rotates as it is wound around a defect center, where a clockwise rotation direction is negative while a counterclockwise rotation is positive. In our system we find $\pm 1/2$ defects using publicly available code to detect them [18]. In addition to these, we also identify $\pm 1$ defects based on the structure of the defect. We use the following steps to detect defects: 1) generate focus-stacked image sequences; 2) calculate the orientation field from these sequences; and 3) detect defects.

Focus stacking fuses the in-focus regions from multiple bright-field images taken at different focal positions of the same region of interest (ROI) into a single image; thus generating an image with larger depth-of-field. We use these to analyze the curved droplet surface. We generate focus-stacked images using a publicly available MATLAB algorithm [19]. Briefly, the algorithm works in three stages: 1) focus measurement; 2) selectivity measure; and 3) image fusion. The regions of focus are measured using gray level local variance, selectivity is estimated from pixel focus to noise ratio, and fusion is performed according to focus measure and features of image pixels categorized by selectivity strength. We obtained focus-stacked image sequences with the following settings: local focus window size = 0.8 – 1.4 $\mu$m; selectivity threshold = 0.4 – 0.7 $\mu$m; and selectivity constant: 0.1 – 0.3.

To calculate the orientation field for each pixel, we use the OrientationJ plugin in ImageJ (v1.53f51) on focus-stacked images. This program generates the orientation field for each pixel using the tensor method, where we set the local structure window size which defines spatial scale for the orientation field calculation. The accuracy of the defects detection mainly depends on the local window size, choosing a smaller window size results in false positives while a larger window size smooths the orientation field, which suppresses defects [20]. We obtain the best results for a window size of 0.5 $\mu$m, which is half the diameter of a bacterium. The structure window size defines the spatial scale over which the orientation field is calculated, and the output is a 32-bit image with an orientation for each pixel with values ranging from $-90°$ to $+90°$. Next, from the orientation field images using MATLAB written code publicly available on GitHub [18], we detect topological defects, their position and direction, and overlaid on the original focus stacked images.

From the orientation field, we calculate the director field by dividing the orientation image into $j \times k$ square grids and obtain vector s by summing the orientations of all pixels in each grid. We use a grid with a 2 $\mu$m diagonal, which is the length of an average cell. To estimate the local nematic ordering, we calculate the nematic order parameter using the equation [21, 18]:

$$Q = \sqrt{\langle \cos 2\theta \rangle^2 + \langle \sin 2\theta \rangle^2},$$

where $\theta$ is the orientation angle and the brackets denote the average for a local window of size 35 $\mu$m$^2$.

#### S4.5 Cell alignment on oil tubes

To calculate the degree of alignment of cells along oil tubes, we calculate the average of the dot products between each cell's orientation vector with the tangent vector of the tube axis in bins of 1.45 $\mu$m. To do this, we first generate a focus-stacked image from confocal slices of an oil tube and then calculate the director for each cell. We then generate the curvilinear axis of the tube that runs along the center, and calculate the unit tangent vector, $\boldsymbol{n}_{\text{tube}}$, in 1.45 $\mu$m width bins. We then calculate $\langle \boldsymbol{n}_{\text{cell},i} \cdot \boldsymbol{n}_{\text{tube}} \rangle$, where $\boldsymbol{n}_{\text{cell},i}$ is the unit vector of the $i$-th cell in bins of 1.45 $\mu$m along the tube axis and the angled brackets represent the average over all cells within the bin.

#### S4.6 Oil tube length measurement

We measure the oil tube length by defining a central axis running from its base to the tip. We assume that functional form for tube elongation is exponential since cell division appears to drive tube elongation and because cells are highly aligned with the central axis of the oil tube (**Fig. 2D,E**). We fit an equation of the form $l_{tube} = L_0 \exp{(t/\tau_{tube})}$ to the mean value of all tube lengths, where $l_{tube}$ is the length of the tube, $L_0$ is the initial tube length, $t$ is time, and $\tau_{tube}$ is the time constant, shown as the dashed line in **Fig. 2F**. We fit both $L_0$ and $\tau_{tube}$. We find that the time required for the tubes to double in length is $\sim$ 3.36 h ($\sim$ 200 min), which is about twice of the cell division time, $t_{div}$, of 1.65 h.

#### S4.7 Pseudo color image generation

We generate the pseudo color images using ImageJ plugin 'Z-stack Depth Colorcode 0.0.2'.

#### S5 Analytical model of oil consumption

We assume that the consumption of oil is mediated by bacteria attached directly to the oil/water interface and the time rate-of-change of the oil volume is:

$$\frac{\mathrm{d}V(t)}{\mathrm{d}t} = -\alpha N(t), \tag{1}$$

where $V(t)$ is the volume of oil, $N(t)$ is the number of interfacial cells, and $\alpha$ is the oil consumption rate of an individual cell (we will define $\alpha_{SB}$ and $\alpha_{DB}$, with subscripts to differentiate the consumption rate in both phenotypes). The maximum number of cells that can lie flat on the interface at a given time is:

$$N(t) = \frac{S(t)}{s_c} c_{\mathrm{pf}} = S(t)\rho, \tag{2}$$

where $S$ is the total surface area at the oil/water interface, $s_c$ is cross sectional area of a cell, and $c_{\mathrm{pf}}$ is the cell packing fraction, which we estimate to be 0.65; $\rho$ is the density of bacteria. We estimate that $s_c = 1.8$ $\mu$m$^2$ for cells 2 $\mu$m in length and 1 $\mu$m in diameter (**Fig. S11B**). Substituting Eq. (1) into Eq. (2), we have:

$$\frac{\mathrm{d}V(t)}{\mathrm{d}t} = -\frac{\alpha c_{\mathrm{pf}}}{s_c} S(t). \tag{3}$$

##### S5.1 Oil consumption by Spherical Biofilms

In the SB phenotype, the oil volume remains approximately spherical as SBs consume the oil. We substitute $4\pi R(t)^3/3$ for $V(t)$ and $4\pi R(t)^2$ for $S(t)$ in Eq. (3), which allows us to relate the time rate-of-change of $R(t)$ to the oil consumption rate of a single cell from the SB phenotype, $\alpha_{\mathrm{SB}}$. Simplifying, we find that:

$$\frac{\mathrm{d}R(t)}{\mathrm{d}t} = -\frac{\alpha_{\mathrm{SB}} c_{\mathrm{pf}}}{s_c} = -\sigma_{\mathrm{SB}}. \tag{4}$$

Integrating Eq. (4), we obtain $R(t) = R_0 - \sigma_{\mathrm{SB}} t$, which we can then normalize by the initial value of the radius ($R_0$) giving:

$$R^*(t) = \frac{R(t)}{R_0} = 1 - \sigma^*_{\mathrm{SB}} t, \tag{5}$$

where $\sigma^*_{\mathrm{SB}} = \sigma_{\mathrm{SB}}/R_0$. This equation agrees with our measurements of drop radii where we see that the time rate-of-change of $R^*(t)$ is a constant, as shown in **Fig. 1D(inset)**. The best fit slope to the $R^*(t)$ data in **Fig. 1D(inset)** yields $\sigma^*_{\mathrm{SB}} = 0.008$ h$^{-1}$. Similar values are obtained for $\sigma^*_{\mathrm{SB}}$ when fitting $S^*$ or $V^*$ in **Fig. 1D,E** through the time evolution functions

$$S^*(t) = (1 - \sigma^*_{\mathrm{SB}} t)^2, \quad \text{and} \quad V^*(t) = (1 - \sigma^*_{\mathrm{SB}} t)^3. \tag{6}$$

From our data of droplet size decrease, we find that $\sigma_{\mathrm{SB}} = 0.26\,\mu\mathrm{m.h}^{-1}$, for $R_0 = 32 \pm 2\,\mu\mathrm{m}$. From such value and Eq. (4), we estimate the average single-cell consumption rate $\alpha_{\mathrm{SB}} = 0.7\,\mathrm{fL.h}^{-1}$. For reference, the volume of a single cell is ~1 fL.

In this model, we assumed that all cells lie flat along the interface, that they all have the same dimensions, and that interfacial cell density $\rho$ is constant.

##### S5.2 Oil consumption by Dendritic Biofilms

Since dendritic biofilms strongly deform the oil volume, we are unable to approximate the decreasing oil volume as a sphere and thus unable to relate $V$ and $S$ to $R$. Although we assume Eqns. (1)-(3) still apply, in this case, we assume that cell division drives the deformations, and therefore that the increase in surface area has an exponential functional form.

From measured division time, $t_{\mathrm{div}}$, shown in **Fig. S1D**, we calculate the exponential time constant as $\tau = t_{\mathrm{div}}/\ln 2$ . To determine the change in $N$ at the oil-water interface, we assume that it scales exponentially with time as:

$$N(t) = N_0 \exp\left(\frac{ft}{\tau}\right), \tag{7}$$

where we include a fitting factor $f$, which represents the fraction of cells at the interface that participate in increasing the interface area. In Sec. S6.2.3, we relate $f$ to the existence of a flux of cells ejected from the interface, into a thick biofilm, which we observe experimentally in **Supplementary Movie 1,2,3,5**. Combining Eq. (2) and (7), we arrive at an equation that relates exponential cell growth to interfacial area:

$$S(t) = \frac{s_c}{c_{\mathrm{pf}}} N_0 \exp\left(\frac{ft}{\tau}\right), \tag{8}$$

which we normalize with the initial value $S_0$, leading to:

$$S^*(t) = \exp\left(\frac{ft}{\tau}\right). \tag{9}$$

We fit the $S^*$ data in **Fig. 1D** using Eq. (9) with $f$ as the only fit parameter, which yields a value of $f = 0.11 \pm 0.01$. According to Eq. (7), this value indicates that $\sim 11\%$ of all cells participate in increasing the surface area, as shown in **Fig. S11A**. Such value is consistent with the homeostatic growth model with oil surface constant elongation rate discussed in S6.2.3). Substituting Eq. (8) into Eq. (3) yields:

$$\frac{\mathrm{d}V}{\mathrm{d}t} = -\alpha_{\mathrm{DB}} N_0 \exp\left(\frac{ft}{\tau}\right) = -\sigma_{\mathrm{DB}} S_0 \exp\left(\frac{ft}{\tau}\right), \tag{10}$$

where $\sigma_{\mathrm{DB}} = \alpha_{\mathrm{DB}} \cdot c_{\mathrm{pf}}/s_c$. Integrating Eq. (10) and utilizing the fact that the droplet is still spherical at $t_0 = 0$, we find that

$$V^*(t) = \frac{V}{V_0} = 1 + 3\sigma^*_{\mathrm{DB}} \frac{t}{\tau} \left[1 - \exp\left(\frac{ft}{\tau}\right)\right]. \tag{11}$$

where $\sigma^*_{\mathrm{DB}} = \sigma_{\mathrm{DB}}/R_0$. Fitting the experimental data with the expression provided in Eq. (11), we find that $\sigma^*_{\mathrm{DB}} = 0.009\ \mathrm{h}^{-1}$. For an $R_0 = 32\ \mu\mathrm{m}$, $\sigma_{\mathrm{DB}} = 0.3\ \mu\mathrm{m}\ \mathrm{h}^{-1}$, and the consumption rate per cell $\alpha_{\mathrm{DB}} = 0.8\,\mathrm{fL.h}^{-1}$. Thus, we find that in the DB phase, although the single-cell consumption rate is $\sim 12\%$ larger than the SB phase, the significant difference in the time rate-of-change of volume in the two phenotypes is rather driven by the exponential growth in interfacial area.

#### S5.3 Surface to Volume Ratio

In the SB phase, the normalized surface-to-volume ratio is

$$\frac{S^*}{V^*} = (1 - \sigma^*_{\mathrm{SB}} t)^{-1}, \tag{12}$$

whereas in the DB phase, it is:

$$\frac{S^*}{V^*} = \frac{\exp\left(\frac{ft}{\tau}\right)}{1 - 3\sigma^*_{\mathrm{DB}} \frac{\tau}{f} \left(\exp\left(\frac{ft}{\tau}\right) - 1\right)}. \tag{13}$$

These equations are used to plot the dashed lines in **Fig. 1E(inset)** using the derived values.

#### S5.4 Data and Statistical Analysis

For data analysis we use MATLABv2020, Origin 2019. For tests of statistical significance, we use Welch's $t$-test.

### S6 Membrane model for spontaneous tube formation

**General free energy** Here, we model the biofilm at the oil/water interface as a membrane with liquid crystal order. The membrane is defined by a metric $g$, a curvature tensor $C$ and a normal vector $\mathbf{e}_n$. The local orientation on the bacteria on the biofilm is described by a director field $\mathbf{n}$, which has nematic symmetry (i.e. $\mathbf{n} \to -\mathbf{n}$) and we consider this orientational order to be well developed (i.e. $|\mathbf{n}| = 1$). We then consider an effective free-energy per unit length

$$\mathcal{F} = \int da \left\{ \gamma + \frac{\kappa_{B,\parallel}}{2} \left(n^i n^j C_{ij} - C_{\parallel,0}\right)^2 + \frac{\kappa_{B,\perp}}{2} \left(n^i_\perp n^j_\perp C_{ij} - C_{\perp,0}\right)^2 + \frac{\kappa_F}{2} (\nabla_i n_j)(\nabla^i n^j) \right\}, \tag{14}$$

where $da$ the element of area along the surface and $\mathbf{n}_\perp = \mathbf{e}_n \times \mathbf{n}/|\mathbf{e}_n \times \mathbf{n}|$ defines the orthogonal direction to the nematic field along the membrane. The first, second and third terms in Eq. (14) account for the energetic cost of deformations of the membrane midplane, where $\gamma$ is the surface tension and $\kappa_{B,\parallel}$ and $\kappa_{B,\perp}$ (resp. $C_{\parallel,0}$ and $C_{\perp,0}$) are the bending rigidities (resp. preferred curvature) of the biofilm in the parallel and perpendicular direction to the local nematic order $\mathbf{n}$ field. The fourth term in Eq. (14) is an energetic cost associated with distortions of the director field with an elastic constant $\kappa_F$.

#### S6.1 Simplified static model

##### S6.1.1 Mechanical stability of a biofilm tube

**Tube geometry** We focus on the mechanical stability of a tube of biofilm. We denote by $z$ the tube axis direction, by $\phi$ the azimuth coordinate and by $r$ the tube radius (**Fig. S14**). We focus on the case of a constant radius tube $r$. For simplicity, we will consider that $C_{0,\parallel} \approx 0$, in which case the second term in Eq. (14) vanishes. Finally, the bacteria main axis of elongation is oriented along the tube $z$ direction, i.e. $\mathbf{n} = (0, 0, 1)$ in the cylindrical coordinate. Because the director field is assumed to be fixed, we disregard energy variations due to distortions of the director fields. In this context, Eq. (14) simplifies into

$$\mathcal{F} = \int \left[ \frac{\kappa_B}{2} \left( \frac{1}{r^2} - \frac{2C_{\theta,0}}{r} \right) + \gamma \right] 2\pi r dz = f_z(r) \int dz. \tag{15}$$

where $C_{\perp,0} = C_{\theta,0}$ is the spontaneous curvature in the $\theta$ direction and where we use the notation $\kappa_B = \kappa_{B,\perp}$; the integral spans over the whole tube surface. In Eq. (15), we defined $f_z$, the free energy per unit length along the $z$ axis. At equilibrium, the normal force balance on the tube [22] reads

$$dr \frac{\partial f_z}{\partial r} = 2\pi r \Delta P dr, \tag{16}$$

where $\Delta P$ is the pressure difference between the inside and outside of the tube. Expressed in terms of Eq. (15), Eq. (16) reads

$$\gamma + \frac{\kappa_B}{2} \left( -\frac{1}{r^2} \right) = r \Delta P. \tag{17}$$

Here, we consider that the pressure within the tube is the same as the pressure within the rest of the oil droplet, which we approximate as a sphere of radius $R_d \gg r$. The pressure difference within spherical part of the oil droplet is $\Delta P = 2\gamma/R_d$ by Laplace law –i.e. Eq. (17) but in a sphere geometry. The right-hand side of Eq. (17) then scales as $r/R_d \ll 1$, which amounts to a small perturbation. Henceforth, we neglect the effect of pressure difference in the rest of the calculation.

The expression of the radius $r_{\text{eq}}$ that satisfies the normal force balance condition Eq. (17) then takes a particularly simple expression:

$$r_{\text{eq}} = \sqrt{\frac{\kappa_B}{2\gamma}}, \tag{18}$$

which is independent of the value of the spontaneous curvature $C_{\theta,0}$. However, the stability of the tube depends on the spontaneous curvature $C_{\theta,0}$. Indeed, the force

$$\frac{d\mathcal{F}}{dz} = f_z, \tag{19}$$

which corresponds to the force needed to pull the tube along the $z$ direction can be expressed in terms of the solution $r_{\text{eq}}$ to Eq. (17) as

$$f_z = 2\pi r_{\text{eq}} \left( 2\gamma - \kappa_B \frac{C_{\theta,0}}{r_{\text{eq}}} \right). \tag{20}$$

The tube shrinks whenever $f_z > 0$ and expands otherwise. The tube stability condition Eq. (20) then reads $f_z = 0$ and is reached for the radius

$$r'_{\text{eq}} = \frac{\kappa_B C_{\theta,0}}{2\gamma}, \tag{21}$$

which, in contrast to Eq. (18), depends on the value of the spontaneous curvature. Based on Eq. (18) and Eq. (21), the condition $r'_{\text{eq}} = r_{\text{eq}}$ yields the threshold spontaneous curvature for tube formation:

$$\gamma = \kappa_B C_{\theta,0}^2/2, \tag{22}$$

which, under the notation $C_{\theta,0} = 1/r_0$, corresponds to Eq. (1) in the main text.

##### S6.1.2 Relation to the bacteria density

We expect the mechanical parameters (bending modulus, spontaneous curvature, and tension) that describe the biofilm to depend on the bacteria density. Here, we consider the expansion at second order in the density field

$$(\gamma - \kappa_B C_{0,\varphi}^2/2)_{|\rho} \sim k_0 - k_1\rho + k_2\rho^2. \tag{23}$$

As the oil/water interface displays no obvious spontaneous curvature in the absence of bacteria ($\rho = 0$), we expect that $k_0 = \gamma_0 > 0$, where $\gamma_0$ is the surface tension in the absence of bacteria. We are then led to define the function

$$\kappa(\rho) = \gamma_0 - k_1\rho + k_2\rho^2. \tag{24}$$

such that the condition of Eq. (22) is met for $\kappa(\rho) = 0$ and that tube formation occurs for $\kappa(\rho) < 0$. We further justify the expansion in $\rho$ in the next section, Sec. S6.2.3. The phenomenology corresponding to Eq. (24) is further discussed in Sec. (S7.3).

#### S6.2 Dynamical model

In this section, we will follow the theoretical framework developed in Ref. [23] and use standard notation of differential geometry, to study the dynamics at the onset of tube formation for an interface made of a growing nematic liquid crystal. For more details on the theoretical framework, we refer to Ref. [23].

##### S6.2.1 Conservation of the bacteria mass

Along the membrane surface, the balance in the density of bacteria $\rho$ can be expressed as:

$$\frac{\partial \rho}{\partial t} + \nabla_i \left(\rho v^i\right) + v_n C_i^i \rho = k(\rho)\rho, \tag{25}$$

where $\nabla_i$ is the covariant derivative, $\mathbf{v} = v^i\mathbf{e}_i + v_n\mathbf{e}_n$ is the local velocity with $\mathbf{e}_i$ basis vectors in the membrane frame of reference and $v_n$ the velocity along the outward-oriented normal vector $\mathbf{e}_n$ to the membrane surface; $k$ is the rate of bacteria turnover, which can be related to the parameter $f$ defined in Eq. (7). Here, we expand $k$ in the vicinity of homeostatic conditions $k(\rho) = (\rho_H - \rho)/(\tau\rho_H)$; for $\rho < \rho_H$, bacteria divide more than they die (or leave the interface), and $k(\rho)$ is a source term; for $\rho > \rho_H$, bacteria are ejected more than they divide, and $k(\rho)$ is a sink term.

In the tube geometry and with $v_n = 0$, Eq. (25) reads

$$\frac{\partial \rho}{\partial t} + v_z \partial_z \rho + \rho \left(\partial_z v^z\right) = \frac{\rho}{\tau}\left(\frac{\rho_H - \rho}{\rho_H}\right). \tag{26}$$

As discussed Sec. (S6.2.2), we will assume a fast mechanical relaxation along the normal direction (i.e. $v_n = \partial r/\partial t = 0$). Here, we neglect spatial heterogeneities along the tube direction, e.g. we consider a uniform bacterial concentration ($\partial_z \rho = 0$) and a constant tube expansion rate $\partial_z v^z$,

$$\partial_z v^z = \frac{1}{L}\frac{dL}{dt}, \tag{27}$$

where $L$ is the total tube length. In this case, Eq. (26) amounts to

$$\frac{d\rho}{dt} + \frac{\rho}{L}\frac{dL}{dt} = \frac{\rho}{\tau}\left(\frac{\rho_H - \rho}{\rho_H}\right). \tag{28}$$

We point out two specific features of Eq. (28):

1. Defining a total number of bacteria $N = \rho(2\pi r L)$, we find that

$$\frac{dN}{dt} = \frac{N}{\tau}\left(\frac{\rho_H - \rho}{\rho_H}\right), \tag{29}$$

   such that the number of bacteria in the biofilm is constant only if $\rho_H = \rho$.

2. In the case of a constant tube growth rate, the steady state ($d\rho/dt = 0$) bacterial density reads

$$\rho_{ss} = \rho_H \left(1 - \frac{\tau}{L}\frac{dL}{dt}\right), \tag{30}$$

   which shows that the steady state density is lower than the homeostatic density due to a dilution effect caused by the tube growth.

##### S6.2.2 Conservation of momentum

**Definition and general force balance equations** Here, considering a surface with a basis of tangent vectors $\mathbf{e}_i$ and normal vector $\mathbf{e}_n$, we recall that, following Ref. [23] notations, the force $\mathbf{f}$ on a line of length $dl$ with unit vector $\boldsymbol{\nu} = \nu^i \mathbf{e}_i$, tangential to the surface of interest and normal to the line can be expressed as

$$\mathbf{f} = dl \nu^i \mathbf{t}_i = dl \nu_i \mathbf{t}^i, \tag{31}$$

where $\mathbf{t}_i$ is a tension that can be expressed into tangential and normal components

$$\mathbf{t}^i = t^{ij}\mathbf{e}_j + t^i_n \mathbf{e}_n. \tag{32}$$

In the absence of inertia, momentum conservation reduces to force balance. In the directions tangent to the surface, force balance can be expressed as

$$\nabla_i t^{ij} + C^j_i t^i_n = (\sigma^{\text{oil}})^j_n - (\sigma^{\text{water}})^j_n. \tag{33}$$

Here, $C_{ij}$ is the curvature tensor and $(\sigma^{\text{oil}})^j_n$ and $(\sigma^{\text{water}})^j_n$ are the normal-$j$ components of the 3d stress tensor in the water and oil phases, respectively. In the direction normal to the surface, force balance reads

$$\nabla_i t^i_n - C_{ij} t^{ij} = (\sigma^{oil})_{nn} - (\sigma^{water})_{nn}. \tag{34}$$

where $(\sigma^{\text{oil}})_{nn}$ and $(\sigma^{\text{water}})_{nn}$ are the normal-normal components of the 3d stress tensor in the water and oil phases, respectively. In the following, we will disregard the effects due to the shear in either the water or the oil interfaced and set the right-hand side of Eq. (33) to zero. In this case, $(\sigma^{oil})_{nn} - (\sigma^{water})_{nn} = \Delta P$, where $\Delta P$ is the pressure difference between the oil and the water phases. As argued in Sec.S6.1, we neglect the effects due to pressure difference within the oil droplet, and set $\Delta P = 0$ from now on.

The constitutive equations of a surface made of a growing nematic liquid crystal constrain the expression for the tension $\mathbf{t}^i$. Here, we consider that the in-plane components $t^{ij}$ take the form

$$t^{ij} = t_e^{ij} - \zeta\Delta\mu g^{ij} - \tilde{\zeta}\Delta\mu(n^i n^j - g^{ij}/2) + \eta_b v_k^k g^{ij} + 2\eta\tilde{v}^{ij}, \tag{35}$$

where $\zeta\Delta\mu$ and $\tilde{\zeta}\Delta\mu$ are active stresses generated by bacteria growth, with $n^i$ the in-plane components of the cell orientation director field; $v_{ij} = (\nabla_i v_j + \nabla_j v_i)/2 + v_n C_{ij}$ is the velocity-gradient tensor and $\tilde{v}^{ij} = v^{ij} - v_k^k g^{ij}/2$ its traceless form. We considered both shear and bulk viscous stresses. The equilibrium stress in Eq. (35) is defined as

$$t_e^{ij} = (f - \mu_b\rho)g^{ij} - K^{ik}C_k^j, \tag{36}$$

where $f$ is a free-energy density, $\mu_b = \partial f/\partial\rho$ is a chemical potential and $K^{ij} = \partial f/\partial C_{ij}$ is the passive bending moment. Furthermore, the normal component of the tension $t_n^i$ takes the form

$$t_n^i = t_{e,n}^i = \nabla_j K^{ji}. \tag{37}$$

**Application to the tube geometry** Considering a cylindrical geometry of radius $r$, which is parametrized as $\mathbf{r} = (r\cos(\theta), r\sin(\theta), z)$, we denote the tangent vectors as: $\mathbf{e}_z = \partial_z\mathbf{r}$ and $\mathbf{e}_\theta = \partial_\theta\mathbf{r}$, and the unit normal vector is $\mathbf{e}_n = \mathbf{e}_\theta \times \mathbf{e}_z/|\mathbf{e}_\theta \times \mathbf{e}_z|$. From these expressions, the metric and curvature tensor are defined as $g_{ij} = e_i \cdot e_j$ and $C_{ij} = -(\partial_{ij}\mathbf{r}) \cdot \mathbf{e}_n$. The non-vanishing components of the previous tensors are: $g_{\theta\theta} = r^2$, $g_{zz} = 1$ and $C_{\theta\theta} = r$. Considering only states that are independent on the azimuthal coordinate $\theta$ and invariant along the axial direction, Eqs. (35–37) take a simpler form. In particular, based on Eq. (14) with $\kappa_B = \kappa_{B,\perp}$, the non-vanishing components of the passive bending moment and the equilibrium in-plane tension read

$$K^{\theta\theta} = \frac{\kappa_B}{r^2}\left(\frac{1}{r} - C_{\theta,0}\right), \tag{38}$$

$$t_e^{zz} = \left(\sigma(\rho) + \frac{\kappa_B}{2r^2} - \frac{\kappa_B C_{\theta,0}}{r}\right), \tag{39}$$

$$t_e^{\theta\theta} = \left(\sigma(\rho) - \frac{\kappa_B}{2r^2}\right)\frac{1}{r^2}, \tag{40}$$

where, considering the possibility of a density-dependent surface tension, we have

$$\sigma(\rho) = \gamma(\rho) - \frac{\partial\gamma(\rho)}{\partial\rho}\rho + \frac{\kappa_B C_{\theta,0}^2}{2}. \tag{41}$$

Note that $t_{e,n}^i = 0$. Besides, the non-vanishing components of the in-plane tension read

$$t^{zz} = \left(\sigma(\rho) - \bar{\zeta}\Delta\mu - \frac{\zeta\Delta\mu}{2} + \frac{\kappa_B}{2r^2} - \frac{\kappa_B, C_{\theta,0}}{r}\right) + (\eta_b + \eta)v^{zz} + (\eta_b - \eta)\frac{v_n}{r}, \tag{42}$$

$$t^{\theta\theta} = \left(\sigma(\rho) - \bar{\zeta}\Delta\mu + \frac{\zeta\Delta\mu}{2} - \frac{\kappa_B}{2r^2} + (\eta_b - \eta)v^{zz} + (\eta_b + \eta)\frac{v_n}{r}\right)\frac{1}{r^2}. \tag{43}$$

Then, the force balance in the axial direction given by Eq. (33) implies that $t^{zz} = t_0^{zz} = cte$, where $t_0^{zz}$ represents an in-plane tension per unit length at the end of the tube. Due to negligible pressure differences between the oil and water phases, this term vanishes (i.e. $t_0^{zz}$). Similarly, force balance in the normal direction given by Eq. (34) implies that $rt^{\theta\theta} = 0$. Therefore, force balance eventually implies that:

$$\sigma_{t,H} + \frac{\kappa_B}{2r^2} - \frac{C_\varphi}{r}\kappa_B + (\eta_b + \tilde{\eta})\frac{\partial v_z}{\partial z} + (\eta_b - \tilde{\eta})\frac{1}{r}\frac{\partial r}{\partial t} = 0, \tag{44}$$

$$\sigma_{n,H} - \frac{\kappa_B}{2r^2} + (\eta_b - \tilde{\eta})\frac{\partial v_z}{\partial z} + (\eta_b + \tilde{\eta})\frac{1}{r}\frac{\partial r}{\partial t} = 0, \tag{45}$$

where we defined

$$\sigma_{t,H} = \sigma(\rho) - \zeta\Delta\mu - \tilde{\zeta}\Delta\mu/2, \quad \text{and} \quad \sigma_{n,H} = \sigma(\rho) - \zeta\Delta\mu + \tilde{\zeta}\Delta\mu/2. \tag{46}$$

**Rapid normal stress equilibration** Here, we will assume that $\eta_b - \tilde{\eta} \ll \eta_b + \tilde{\eta}$, and that the tube radius remains constant at steady state. This amounts to neglecting all dynamical variables in Eq. (45) resulting in the condition:

$$\sigma_{n,H} - \frac{\kappa_B}{2r_{\text{eq}}^2} = 0, \tag{47}$$

which sets the tube radius at

$$r_{\text{eq}} = \sqrt{\frac{\kappa_B}{2\sigma_{n,H}}}. \tag{48}$$

In this limit, Eq. (44) takes the expression:

$$\eta \frac{1}{L}\frac{dL}{dt} = -\left(\sigma_{t,H} + \sigma_{n,H} - \frac{C_{\theta,0}}{r_{\text{eq}}}\kappa_B\right), \tag{49}$$

where $\eta = \eta_b + \tilde{\eta}$; $dL/dt$ is defined in Eq. (27). With all parameters being fixed, Eq. (49) predicts an exponential tube growth at a rate $-\kappa/\eta$, with

$$\kappa \equiv \sigma_{t,H} + \sigma_{n,H} - \frac{C_{\theta,0}}{r_{\text{eq}}}\kappa_B, \tag{50}$$

which corresponds to the Eq. (1) provided in the main text, and, in the case $\sigma_{t,H} = \sigma_{n,H} = \gamma$, to Eq. (20). Tube formation then occurs when $\kappa < 0$.

**Experimental estimates** We consider a bending energy of the order of $\kappa_B \approx Ew^3 = 10^{-12}\,\text{N.m}$, with $E = 10^6\,\text{Pa}$ and an estimated bacteria biofilm thickness $w = 10^{-6}\,$ m. an equilibrium tube radius of the form $r_{\text{eq}} = 10^{-5}\,\text{m}$ and a preferred radius $r_0 = 5 \cdot 10^{-6}\,\text{m}$. This leads to a negative contribution to the overall effective surface tension $\kappa_{\text{act}} \sim \kappa_B/(r_0 r_{\text{eq}}) \sim 20\,\text{mN.m}$, which is in the range of the measured surface tension values. The tube growth rate $\tau \sim \eta/\kappa$; $\tau \approx 350\,\text{min} \approx 10^4\,\text{s}$ in experiments. Considering a two-dimensional biofilm layer viscosity $\eta = 1\,\text{N.m}^{-1}$.s (constructed as $\eta = \eta_{3\text{D}} w$, with $\eta_{3\text{D}} = 10^6\,\text{N.m}^{-2}$.s and $w = 10^{-6}\text{m}$) implies that the total effective surface tension needs to be relatively small, in the $\kappa \sim 0.1\,\text{mN.m}$ range.

**Condition for tube formation** Inserting Eq. (47) into Eq. (49) for $dL/dt = 0$ yields the following critical equilibrium radius:

$$r'_{\text{eq}} = \frac{\kappa_B C_{\theta,0}}{\sigma_{t,H} + \sigma_{n,H}}. \tag{51}$$

The condition $r'_{\text{eq}} = r_{\text{eq}}$ yields the critical surface tension condition:

$$\frac{(\sigma_{t,H} + \sigma_{n,H})^2}{2\sigma_{n,H}} < \kappa_B C_{\theta,0}^2. \tag{52}$$

In all the equations defined here in Sec. S6.2.2, there are no reference to the bacteria density field. This will be the subject of the next paragraph.

##### S6.2.3 Coupling the stress equation to the bacterial density: oscillation analysis

Here, we show that the transition between the spherical and dendritic phenotypes – as well as the existence of oscillations between these phenotypes – is recapitulated by through a relation between the tube growth rate and the density of bacteria in the biofilm. We focus on the limit of a fast division rate $1 \ll \kappa\tau/\eta$, such that $\rho \approx \rho_{ss}(\dot{L})$, where $\rho_{ss} = 1 - \tau\dot{\epsilon}$ is the steady-state density defined in Eq. (30), with $\dot{\epsilon} = \dot{L}/L$. The tube growth rate defined in Eqs. (49) and (50) can then be expressed as:

$$\eta\dot{\epsilon} = -\kappa(\rho), \tag{53}$$

where we now assume a functional coupling between $\kappa$ and $\rho$; we approximate such coupling to its second order expansion around $\rho_{ss}$:

$$\kappa \sim k_0 - k_1\rho + k_2\rho^2. \tag{54}$$

Most likely the expansion in $\rho$ in Eq. (54) specifically originates from the active stress contribution $\sigma_{t,H}$ in Eq. (49). A similar third-order expansion of such term was considered in Ref. [24].

**Relation to the growth rate correction $f$, Eq. (7)** In the case of a constant tube elongation rate $\dot{\epsilon}$ (dendritic growth), the density $\rho_{ss}$ is constant; in this case, Eq. (29) yields an expression for the total number of bacteria that is consistent with Eq. (7) with $f = k_d(1 - \rho_{ss}/\rho_H) > 0$.

**Oscillations** Expanding Eq. (54) at first orders in the tube growth rate $\dot{\epsilon}$ then leads to

$$\kappa \sim \hat{k}_0 - \hat{k}_1 \tau \dot{\epsilon} + \hat{k}_2 \tau^2 \dot{\epsilon}^2. \tag{55}$$

where $\hat{k}_0 = k_0 - k_1\rho_H + k_2\rho_H^2$, $\hat{k}_1 = k_1 - 2k_2\rho_H$, $\hat{k}_2 = k_2 > 0$. Given Eq. (55), there can be two stable solutions to Eq. (53), one corresponding to a collapsing velocity $\dot{\epsilon}_c < 0$ and the other corresponding to a growth velocity $\dot{\epsilon}_g > 0$. Processes limiting the growth or collapse of the biofilm length can trigger a switch between these two stable solutions. The transition from the growth phase to the collapse phase can be triggered when the tube length reaches a maximum, as for example when the disappearance of the oil droplet leads to an increase of tube oil pressure. In turn, the collapse phase eventually stops when the total interface length reaches the minimal critical value set by the quantity of oil within the droplet, which leads to the next growth phase. The mechanism we propose here is reminiscent of the one proposed for the periodic cycles observed in molecular motors assemblies [25].

In the next section, we propose a phase-field model where oscillations in the modelled tube length emerge spontaneously for an intermediate range of parameters.

### S7 Phase-field model for the spherical-dendritic oscillations

#### S7.1 Shape of the oil droplet: a phase field model

Here, we describe a phase field model to describe the shape of the oil droplet, where the field $\phi$ models the local relative fraction in oil. For simplicity, we focus on the two-dimensional configuration with periodic boundary conditions and we consider an initially circular droplet of radius $R$, with $\phi = 1$ within the oil droplet $r < R$ and $\phi = 0$ elsewhere (see Sec. S7.5 for more details).

**Model description** In the absence of oil consumption, we consider a Cahn-Hilliard-like equation for the oil fraction field in the form:

$$\frac{\partial \phi}{\partial t} = M \nabla^2 \mu_\phi, \tag{56}$$

with a chemical potential

$$\mu_\phi = \frac{\delta \mathcal{F}_{CH}}{\delta \phi}, \tag{57}$$

defined in terms of a Canham-Helfrich free energy

$$\mathcal{F}_{CH} = \int \mathrm{d}^2\mathbf{r} \left\{ \frac{1}{2} a \phi^2 (1-\phi)^2 + \frac{1}{2}\kappa_1 (\nabla \phi)^2 + \frac{1}{2}\kappa_2 \left(\nabla^2 \phi\right)^2 \right\}. \tag{58}$$

In two-dimension, the term $\kappa_1$ corresponds to an effective surface tension; $\kappa_2$ corresponds to a Helfrich energy; the ratio of the parameters $a$ and $\kappa_2$ control the width of the oil/water interface, which scales as $\ell \sim (\kappa_2/a)^{1/4}$. We choose a set of parameters such that the length of the interface remains sharp.

**Model results** We first simulated the behavior of Eq. (56) varying the value of $-\kappa_1$ with all other parameters fixed at the values defined in Sec. S7.5. see As illustrated S15, we find that digitation (representing tubes) are observed only for sufficiently negative values of the effective surface tension

$$\kappa_1 < \kappa_{c,1} \approx -0.75. \tag{59}$$

#### S7.2 Bacterial population at the oil-water interface: a logistic growth model

We now model the bacterial population growth. At low concentrations, bacteria grow exponentially until reaching a saturation threshold, which we denote $\rho_H$. A standard model for such population dynamics is the logistic growth [26], where the number of bacteria within the interface (denoted $N$) reads

$$\frac{dN}{dt} = k_d(1 - \rho/\rho_H)N. \tag{60}$$

Dividing by the length of the interface, we obtain the following expression for the density of bacteria

$$\rho(t) = N(t)/\mathscr{L}(t), \tag{61}$$

where, following Ref. [27], the total oil-droplet interface length $\mathscr{L}(t)$ is defined as

$$\mathscr{L}(t) = \int \mathrm{d}^2\mathbf{r}|\nabla\phi|. \tag{62}$$

#### S7.3 Modulation of the oil surface tension by the bacteria density

Inspired by Eqs. (24) and (54), we consider the interfacial term defined in Eq. (58) to read:

$$\kappa_1(\rho) = k_0 - k_1\rho + k_2\rho^2 = k_2(\rho - \rho_1)(\rho - \rho_2). \tag{63}$$

Solving for $\kappa_1(\rho_1) = \kappa_{c,1}$, as defined in Eq. (59), yields two critical densities

$$\rho_1 = \frac{k_1 - \sqrt{k_1^2 - 4\tilde{k}_0 k_2}}{2k_2}, \qquad \text{and} \qquad \rho_2 = \frac{\sqrt{k_1^2 - 4\tilde{k}_0 k_2} + k_1}{2k_2}, \tag{64}$$

with $\tilde{k}_0 = k_0 - \kappa_{c,1}$; these are positive when $k_1^2 - 4\tilde{k}_0 k_2 > 0$ and $k_1, k_2 > 0$. As discussed in Sec. S7.1 and Fig. S15, tube form for any value constant value of the density $\rho \in (\rho_1, \rho_2)$, $\kappa_1(\rho_1) < \kappa_{c,1}$. We then turn to simulations where Eq. (60) is implemented together with the coupling Eq. (63). Different regimes appear according to the value homeostatic density $\rho_H$ with respect to $\rho_1$, $\rho_2$ and the density minimizing $\kappa_1(\rho)$, denoted as $\rho_S$. We consider a spherical droplet with $\rho = 0.001$ at $t = 0$. Such droplet

- (for $\rho_1 < \rho_H < \rho_S = k_2/k_1 < \rho_2$) evolves into a stable dendritic structure, see **Supplementary Movie 4**. This behavior mimics the dendritic phenotype observed for long culture time.
- (for $\rho_1 < \rho_S = k_2/k_1 < \rho_H < \rho_2$) exhibits sustained oscillations between a spherical and a dendritic structure, see SI Fig. S16 and **Supplementary Movie 4**. This behavior mimics the oscillatory behavior observed in experiments at intermediate culture time.
- (for $\rho_H < \rho_1$) remain spherical, see **Supplementary Movie 4**. This behavior mimics the spherical phenotype observed for short culture time.
- (for $\rho_H > \rho_2$) first evolves into a dendritic structure that eventually collapses back into a sphere.

This set of behavior is recapitulated in the phase diagram of SI Fig. S17.

#### S7.4 Relating the phase-field model to the membrane model of Sec. S6.2

**Relating $\kappa_1$ (phase-field model) to $\kappa$ (membrane model)** Tube form in the phase-field model whenever $\kappa_1$ is lower than a threshold value $\kappa_1 < \kappa_{c,1}$, see Fig. S15. We identify the condition $\kappa_1 < \kappa_{c,1}$ to the condition $\kappa < 0$ set in Eq. (53). As shown in Fig. S15, in the range $\kappa_{c,1} < \kappa_1 \in (-0.8, -0.95)$, the strain-rate $\dot{\mathscr{L}}/\mathscr{L}$ increases linearly with $-\kappa_1$ thus mimicking the behavior of $\dot{L}/L$ described in Eq. (53).

**Relating $M$ (phase-field model) to $\eta$ (membrane model)** The time-scale unit in Eq. (58) is set by the parameter $M$ while, in the continuum membrane model Eq. (35), it is set by $\eta_b + \tilde{\eta}$.

**Relating the growth rate $k_d$ (phase-field model) to $\tau$ (membrane model)** Expressing the time derivative of Eq. (61) in terms of Eq. (60), we obtain a relation analogous to Eq. (28), with $k_d = 1/(\rho_H \tau)$.

**Relating $\kappa_2$ (phase-field model) to $\kappa_B$ (membrane model)** Once in the dendritic phenotype, we find that fingers are of thickness $2\ell \sim (\kappa_2/a)^{1/4}$; we therefore map the interface thickness to the tube radius defined Eq. (48), $\ell \sim r_{\text{eq}} = \sqrt{\kappa_B/(2\sigma_{n,H})}$.

##### S7.5 Numerical implementation

We employ a finite difference method to solve the governing equations (Eqs. (56) and (60)). The time integration is performed using a forward Euler scheme; the spatial derivatives are carried out using a second order central difference. Simulations were performed on a 256 × 256 two-dimensional lattice using periodic boundary conditions. The system was initialized with a circular shape of oil of radius $R$ centered within the simulation domain, that is $\phi = 1$ for $r = \sqrt{x^2 + y^2} \leq R$; otherwise $\phi = 0$. If not considered otherwise, we consider the following parameter values: $a = 10$, $M = 0.5$, $\kappa_2 = 0.1$, $k_d = 0.1$, $k_0 = 1.35$, $k_2 = 3.0$, $R = 5$, $\Delta x = 0.125$, $\Delta t = 0.001$. The other parameter values (which are recalled in **Supplementary Movie 4** caption) read

- for the SB phenotype: $k_1 = 5.0$, $\rho_H = 1.2$.
- for the OB phenotype: $k_1 = 5.4$, $\rho_H = 1.2$.
- for the DB phenotype: $k_1 = 5.4$, $\rho_H = 1$.

##### S7.6 Model extensions: including oil consumption

**Including fluctuations** Fluctuations in the droplet shape can be considered through a conservative fluctuating current term, denoted $\vec{J}$

$$\frac{\partial \phi}{\partial t} = M \nabla \cdot \left[ \nabla \mu_\phi + \vec{J} \right], \tag{65}$$

where $\vec{J}$ is a fluctuating Gaussian white noise vector, with zero mean and correlations $\langle J_i(\boldsymbol{x}, t) J_j(\boldsymbol{x}', t') \rangle = \Sigma^2 \delta(\boldsymbol{x} - \boldsymbol{x}') \delta(t - t') \delta_{ij}$. In the **Supplementary Movie S4**, we considered a simulation with $\Sigma = 0.05$. Adding such noise allows for more realistic simulations.

**Including oil consumption** We have not considered the effect of oil consumption by the biofilm. The quantity $\mu_\phi$ defined in Eq. (57) is indeed a Lagrange multiplier that conserves the integrated value of $\phi$. Oil consumption by the biofilm could be investigated by considering the following expression for the oil droplet evolution equation:

$$\frac{\partial \phi}{\partial t} = M \nabla^2 \mu_\phi - k_c |\nabla \phi|, \tag{66}$$

such that $k_c > 0$ corresponds to the rate of oil consumption. We leave the impact of oil consumption to further studies.

#### S8 Fit of the topological-defects mediated dimple profiles

In the following, we detail the procedure for estimating the parameters of a theoretical description of a membrane with nematic order from experimental profiles of dimples near aster topological defects on flattened biofilms.

##### S8.1 Derivation of the fitting formula

Here, we provide a derivation of the linearized shape equation based on Ref. [28]. For more details on the theoretical framework and derivation of the nonlinear shape equation, we refer to Refs. [28, 29].

**Disk geometry** To model the experimental observation of dimple formation near aster topological defects (as described in Fig. 4 in the main text), we consider a flat circular domain of radius $R_{\rm dimple}$ (representing the edge of the visible droplet). Following the observation of Fig. S10, the bacterial orientation is represented by a fixed director field $\mathbf{n}$ along the radial direction $\mathbf{e}_r$ and with modulus $|\mathbf{n}|=1$. The biofilm surface is parameterized by the position vector $\mathbf{r}=(r\cos(\theta), r\sin(\theta), h(r))$, where $h(r)/R_{\rm dimple}\ll 1$ and $h'(r)\ll 1$. The corresponding basis of tangent vectors are: $\mathbf{e}_r=\partial_r\mathbf{r}=(\cos(\theta),\sin(\theta),h'(r))$ and $\mathbf{e}_\theta=r(-\sin(\theta),\cos(\theta),0)$ and $\mathbf{e}_n=(-h'(r)\cos(\theta),-h'(r)\sin(\theta),1)/\sqrt{1+h'(r)^2}$ is the unit vector normal to the surface. With these notations, the elementary surface area defined in Eq. (14) reads $da=drd\theta\sqrt{\det(g_{ij})}$ where $\det(g_{ij})$ is the determinant of the metric tensor. The non-vanishing components of the metric tensor $g_{ij}=\mathbf{e}_i\cdot\mathbf{e}_j$ are $g_{rr}=1+h'(r)^2$ and $g_{\theta\theta}=r^2$, leading to $\det(g_{ij})=(1+h'(r)^2)r^2$. The non-vanishing components of the curvature tensor $C_{ij}=-(\partial_{ij}\mathbf{r})\cdot\mathbf{e}_n$ are $C_{rr}=-h''(r)/\sqrt{1+h'(r)^2}$ and $C_{\theta\theta}=-rh'(r)/\sqrt{1+h'(r)^2}$. This leads to the expression of the mean curvature

$$C_\perp^2+C_\parallel^2\sim\left(-h''(r)-\frac{1}{r^2}\times rh'(r)\right)^2, \tag{67}$$

where we have used that $g_{rr}\sim 1$ at leading order in $h'\ll 1$.

**Result** We consider the free energy of Eq. (14) with $\kappa_{B,\parallel}=\kappa_{B,\perp}$ and ignore boundary terms. We set for simplicity that $C_{0,\perp}=C_{0,\parallel}=0$; this amounts to supposing that at the early stage of the dimple formation considered here, the effect of the spontaneous curvature $C_0$ is not apparent. In this context and using Eq. (67), at leading order in the perturbations $h/R_{\rm dimple}, h'\ll 1$ the free energy expression Eq. (14) reduces to

$$\mathcal{F}\approx\int drd\theta\left\{\gamma r\left(1+\frac{h'(r)^2}{2}\right)+\frac{\kappa_B}{4}r\left(h''+\frac{h'}{r}\right)^2+\frac{\kappa_F}{2r}\left(1-\frac{h'(r)^2}{2}\right)\right\}. \tag{68}$$

The minimal steady-state $h(r)$ profiles satisfy the condition $\delta\mathcal{F}/\delta h=0$, which reads as

$$-\frac{\gamma}{r}\partial_r\left(r\partial_r h\right)+\frac{\kappa_B}{2r}\partial_r\left(r\partial_r(\frac{1}{r}\partial_r\left(r\partial_r h\right)\right)+\frac{\kappa_F}{2r}\partial_r\left(\frac{\partial_r h}{r}\right)=0. \tag{69}$$

This equation is, to linear order, the shape equation of a membrane with nematic order and an aster topological defect at $r=0$ [28]. Integrating Eq. (69) with respect to the radial coordinate $r$ and enforcing the integration constant to vanish, which corresponds to the case of a free boundary at $r=R_{\rm dimple}$, one obtains:

$$-\gamma r\partial_r h+\frac{\kappa_B}{2}\left(r\partial_r(\frac{1}{r}\partial_r\left(r\partial_r h\right)\right)+\frac{\kappa_F}{2}\left(\frac{\partial_r h}{r}\right)=0. \tag{70}$$

The latter equation has a simple solution:

$$\partial_r h=A_1 I\left(\sqrt{1-\frac{\kappa_F}{\kappa_B}},\frac{r}{\sqrt{\kappa_B/\gamma}}\right)+A_2 K\left(\sqrt{1-\frac{\kappa_F}{\kappa_B}},\frac{r}{\sqrt{\kappa_B/\gamma}}\right), \tag{71}$$

where $I(\alpha,x)$ and $K(\alpha,x)$ are Modified Bessel Functions of order $\alpha$ and both $A_1$ and $A_2$ are integration constants. Because, $I(\alpha,x)$ diverges when $x\to\infty$, we set $A_1=0$, and thus the minimal profiles $h(r)$ take the form

$$h=A_2\int_{R_{\rm dimple}}^{r}K\left(\sqrt{1-\frac{\kappa_F}{\kappa_B}},\frac{r'}{\sqrt{\kappa_B/\gamma}}\right)dr'=A_2R_{\rm dimple}\int_1^{r/R_{\rm dimple}}K\left(\sqrt{1-\frac{\kappa_F}{\kappa_B}},\frac{z}{\sqrt{\kappa_B/\gamma R_{\rm dimple}^2}}\right)dz, \tag{72}$$

where $z=r/R_{\rm dimple}$. The last integration constant corresponds to a vertical translation and we fix it to satisfy $h(r=R_{\rm dimple})=0$. The radius $R_{\rm dimple}$ sets the system radius and in experiments, this corresponds to the typical size of flattened oil droplets. As a fitting function we use

$$h^{\rm fit}(r)=(h_0/N)\int_1^{r/R_{\rm dimple}}K\left(\sqrt{1-\frac{\alpha_2}{\alpha_1}},\frac{z}{\sqrt{\alpha_1}}\right)dz, \tag{73}$$

where the fitting parameters are $h_0$, $\alpha_1$ and $\alpha_2$, with the latter parameters defined as

$$\alpha_1 = \kappa_B/\gamma R^2_{\text{dimple}}, \quad \text{and} \quad \alpha_2 = \kappa_F/\gamma R^2_{\text{dimple}}. \tag{74}$$

and $N$ is a normalization factor, defined as

$$N = \int_1^{r_{min}/R_{\text{dimple}}} K\left(\sqrt{1-\frac{\alpha_2}{\alpha_1}}, \frac{z}{\sqrt{\alpha_1}}\right) dz. \tag{75}$$

with $r_{min}/R_{\text{dimple}} = 0.005$.

**Discussion** In Ref. [28], Eq. (69) was generalized for a membrane with an active nematic liquid crystal, and the authors found that the anisotropic active stresses renormalized the surface tension $\gamma$ and the elastic constant $\kappa_F$ in a space-dependent manner. In the experiments considered here, we do not observe significant motility of the defects during the dimple formation process, hence we neglected the effects of anisotropic active stresses to explain the dimple formation.

#### S8.2 Fitting procedure

For a given set of the fitting parameters $(h_0, \alpha_1, \alpha_2)$, we compute the error function

$$\mathcal{E} = \sqrt{\sum_{n=1}^{4}\sum_{r_i} |h^{\exp}(r_i/R_{\text{dimple}}) - h^{\text{fit}}(r = r_i/R_{\text{dimple}})|^2}, \tag{76}$$

where $h^{\exp}(r_i)$ corresponds to the experimental value of the dimple height at the radial position $r = r_i$ normalized by the radius $R_{\text{dimple}}$. The experimental profiles $h^{\exp}(r_i)$ are normalized so that the height at the first position near the peak, which is typically at $r_{\text{min}}/R_{\text{dimple}} = 0.005$, is $1$. The radial position $r_i$ was computed with respect to the position of the maximal peak. The sum in Eq. (76) runs over all experimental position $r_i$ at a given instant of time and four experimental profiles $n = 4$. The latter experimental profiles correspond to two profiles on two perpendicular planes that cross the center of the oil droplet. The analysis was performed over $N = 6$ independent oil droplets. For a given experiment, profiles at different time points are grouped by the value of their maximal height. For this fitting analysis, the typical radial step is $\Delta r = 1.3$ $\mu$m, which is ten times larger than the experimental resolution. The offset of the experimental dimple height was the averaged height at large distances from the position of the maximal peak. For the data set that was analyzed, the dimple radius ranges from $R_{\text{dimple}} = 25-35\mu$m and the height of dimples ranges from $4-8$ $\mu$m.

The error function $\mathcal{E}$ was computed in the parameter space $(h_0, \alpha_1, \alpha_2) = (10^{-1}, 10) \times (1, 10^4) \times (1, 10^4)$. We searched for the absolute minimum $\mathcal{E}_{min}$ of the error function $\mathcal{E}$ over the parameter space $(h_0, \alpha_1, \alpha_2)$. Fig. S12 shows the subset of the previous parameter space whereby the error function $\mathcal{E} < 1.2 * \mathcal{E}_{min}$. Our analysis disclosed regions of the parameter space that are compatible with the experimental measurements, Fig. S12. Fig. S13 shows the averaged theoretical fits for each independent experiment. To compute the fitting parameters from Table S2, we used the regions of parameters from Fig. S12 and found that the ratio $\kappa_F/\kappa_B$ is bounded in most of the experimental cases to a value below $5.5$ and in some cases it is constrained to an average value between $1.7 - 2.0$. In all experimental cases, the averaged value of $h_0$ is close to $1$. In one experimental case, we compared experimental profiles with a height $< 6$ $\mu$m and $> 6$ $\mu$m (Fig. S13) and found no differences in the parameter values $\kappa_B/\gamma R^2_{\text{dimple}}$ and $\kappa_F/\gamma R^2_{\text{dimple}}$ (Fig. S12a and Table S2).

### S9 Supplementary Figures

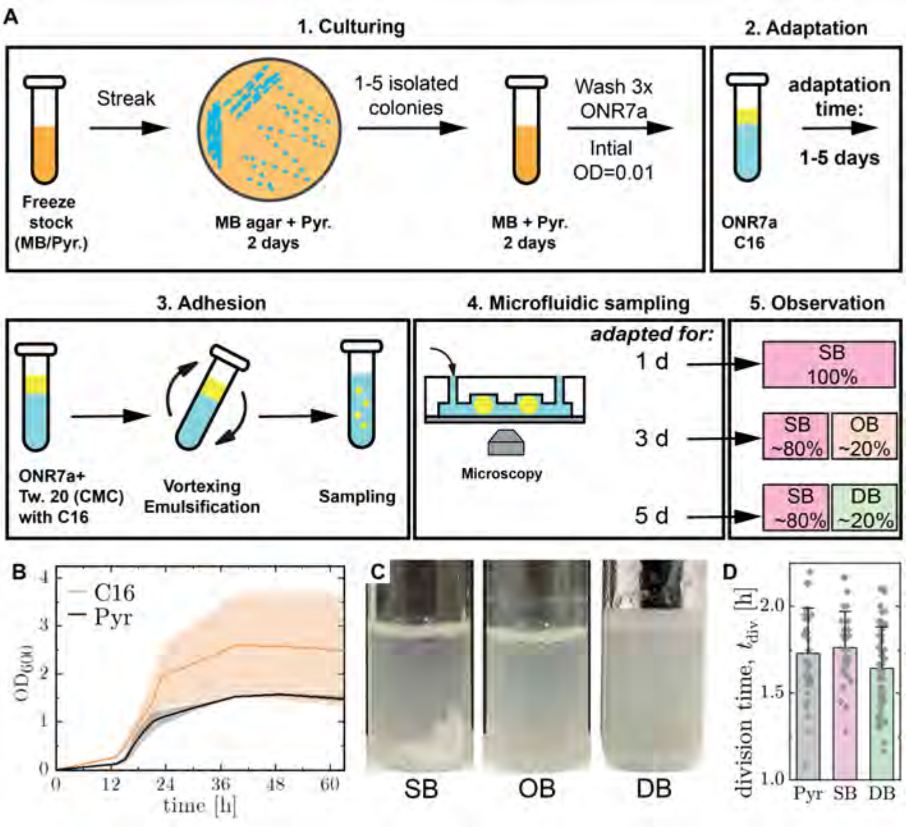

**Figure S1:** (A) Diagram of the: 1) culture process; 2) adaptation of *A. borkumensis* to growth on C16; 3) adhesion to fresh C16 and drop formation; 4) microfluidic sampling; and 5) isolation of SB, OB, and DB phenotypes based on microfluidics-based identification. (B) Growth curves of *A. borkumensis* grown in ONR7A media supplemented with hexadecane (C16) or pyruvate (Pyr), respectively. (C) Images of test tube cultures taken after 5 d of growth. The SB phenotype develops a clearly visible biofilm near the bottom of the test tube attached to the wall. The OB lacks any visible biofilm but large emulsion droplets are present. The DB phenotype is more turbid than the other two, also lacks visible biofilm attached to the tube, and has emulsions much smaller in size. (D) The average division time ($t_{\mathrm{div}}$) of cells grown on Pyr (n=37), SB-forming cells grown on C16 (n=36 cells), and DB-forming cells grown on C16 (n=61 cells), respectively.

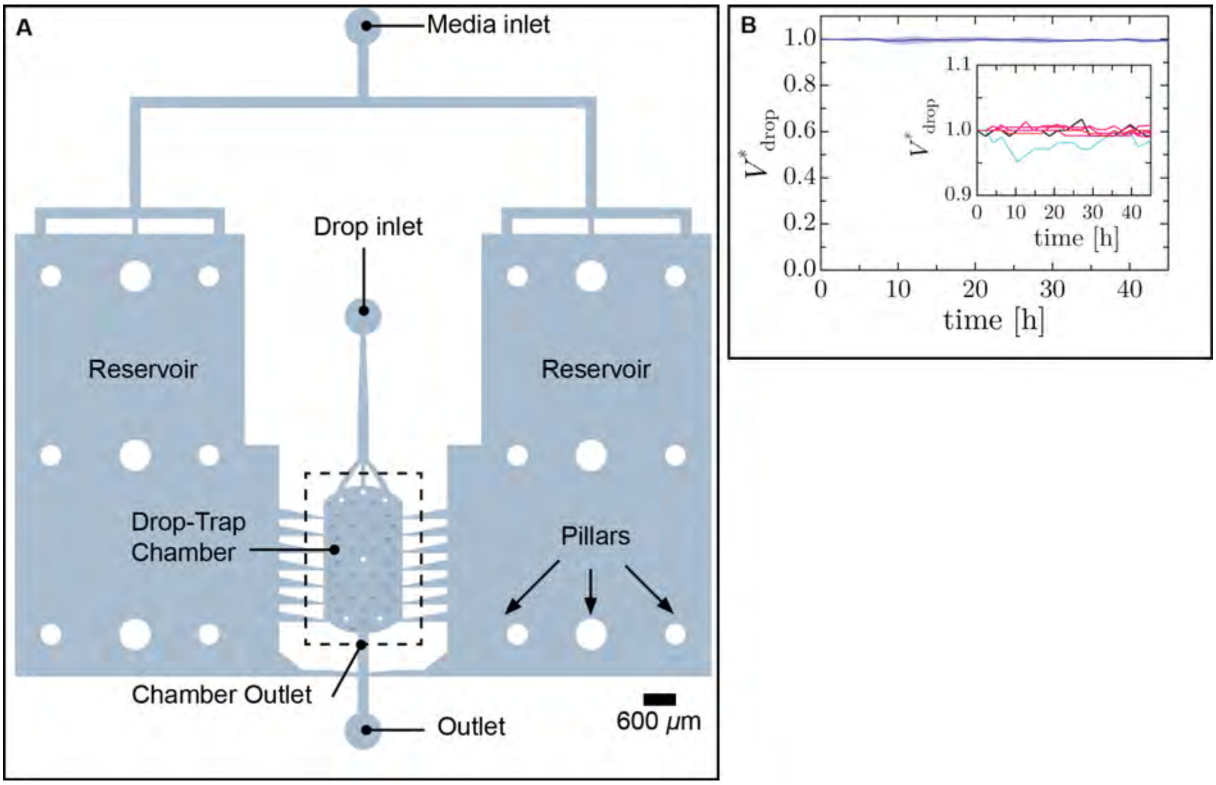

**Figure S2:** (A) Schematic of the 2-layer microfluidic oil-drop trap device used in these experiments. The media-filled channels are shown. The dashed line indicates the drop-trap chamber, which is shown in **Fig. 1A**. The white circles are pillars and small 'raised' blue circles are the drop traps. The media inlet connects to reservoirs that provide a gentle flow through the trap chamber. The drop inlet port is used only to introduce the cell-laden drops into the device. (B) Normalized volume of trapped oil drops over time. Devices are coated with PVA to prevent absorption of C16 by the PDMS channels, which have been measured to suppress absorption for at least 21 days. The solid line and filled area represent the mean ± s.d. (n = 15). (inset) Magnified view of the individual experiments.

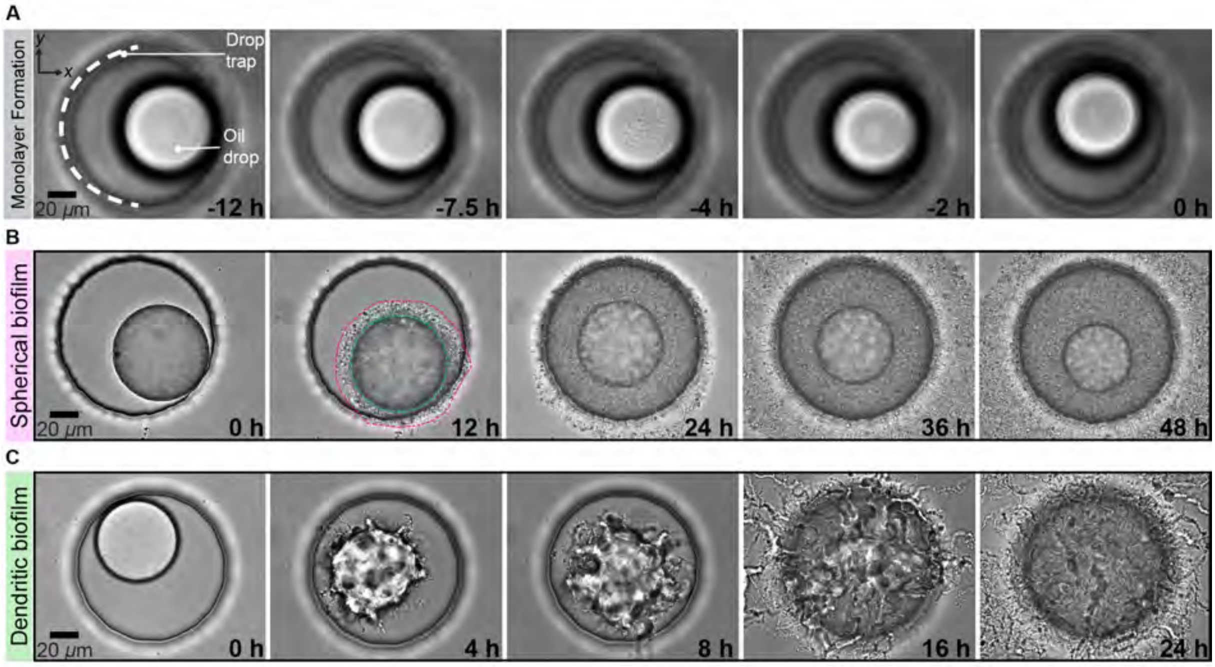

**Figure S3:** (A) Time-lapse sequence showing the formation of a monolayer on a trapped drop. The edge of the drop trap is indicated by the white dashed line. The confluent monolayer is formed at $t_0$ (t = 0 h). (B) Development of a spherical biofilm on an oil drop. The oil drop radius monotonically decreases in time. At 12 h, the droplet is outlined with a cyan line and the biofilm is outlined by a magenta line, as a guide for the reader. (C) Time-lapse sequence showing the development of a dendritic biofilm on an oil drop. The biofilm deforms the surface, initially generating wrinkles and protrusions, that fragmenting the droplet into tube-like segments and numerous smaller irregularly shaped volumes of oil covered with cells at later times.

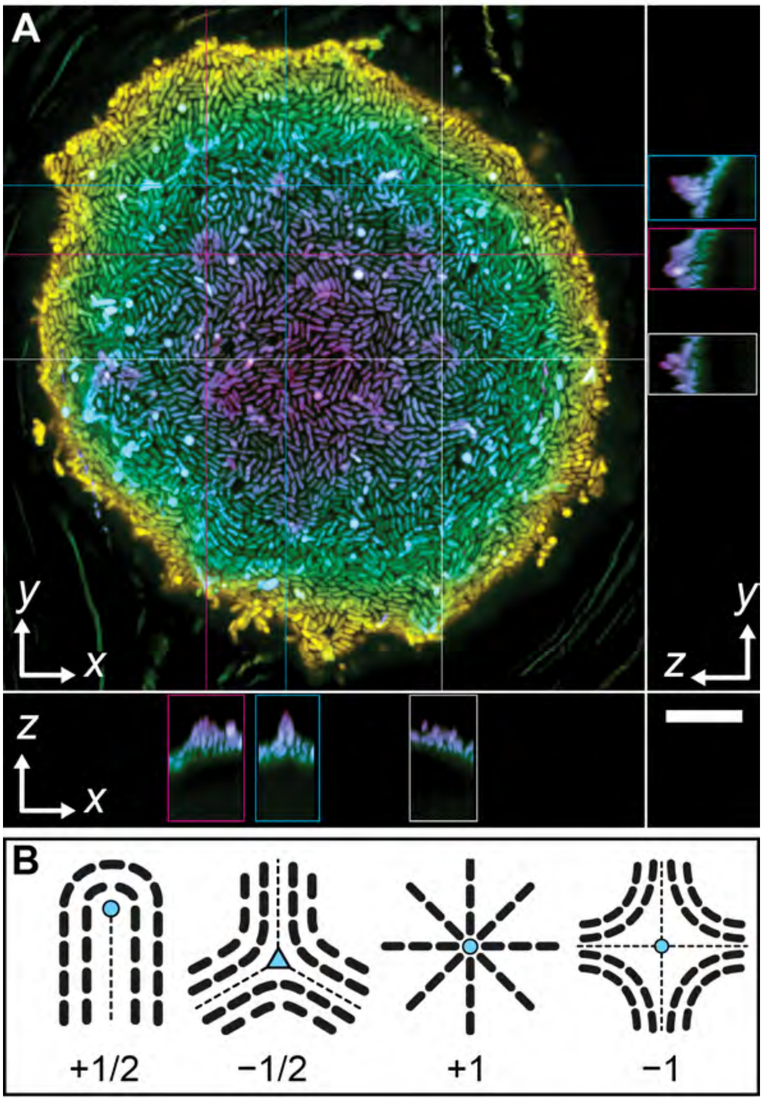

**Figure S4:** Maximum intensity projection confocal image of a droplet at $\sim 10$ h into biofilm development. The colored cross-hairs intersect different $+1$ topological defects, shown in the orthogonal planes, below and to the right. Scale bar $= 10$ $\mu$m. The image is color coded by depth with the colormap scale ranging from 0-19 $\mu$m. (B) Schematic of nematic defects with charges of $\pm 1/2$ and $\pm 1$.

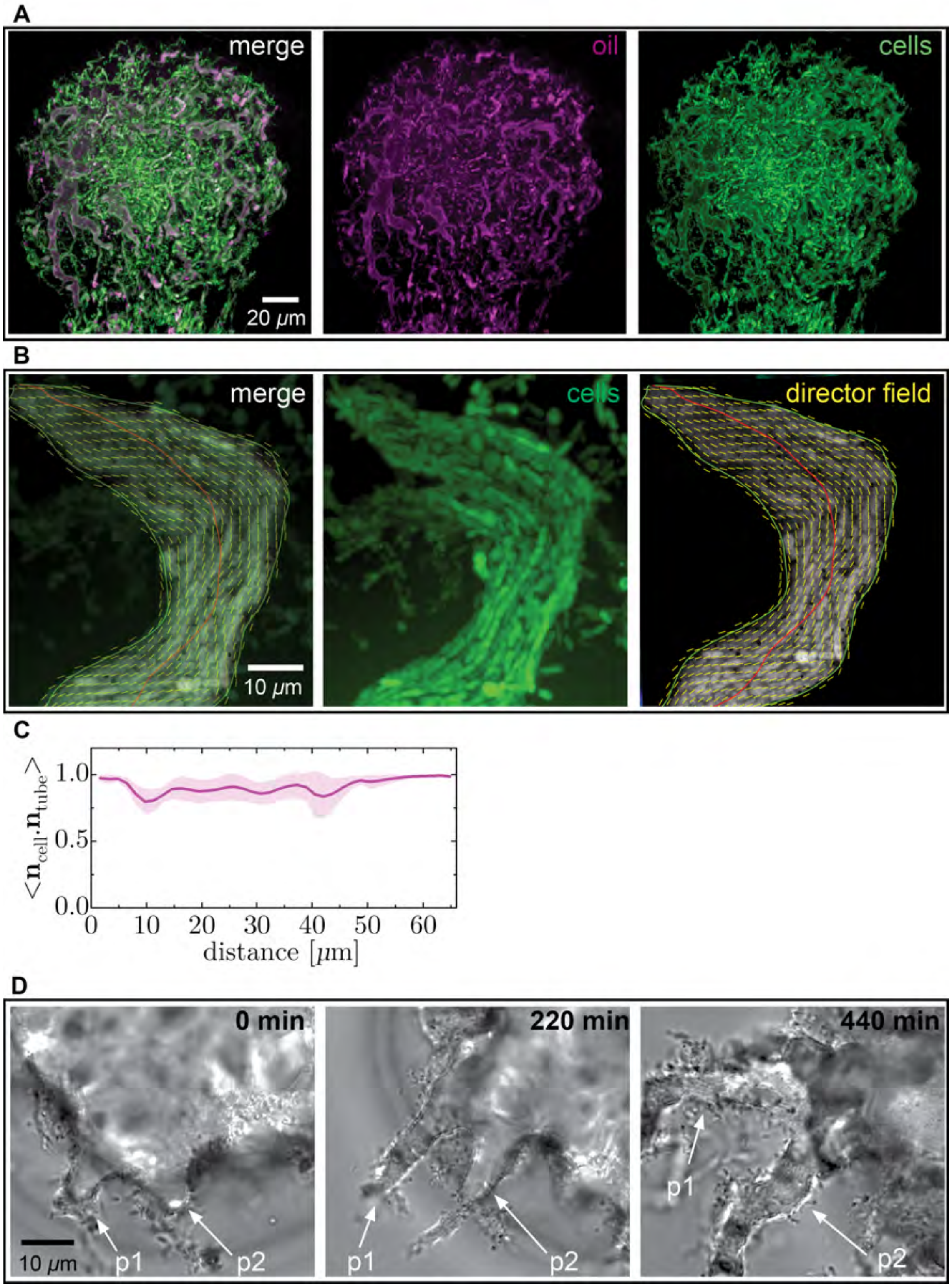

**Figure S5:** (A) A merged 2-color confocal image of an oil drop in the later stages ($\sim$ 15 h) of deformation by a DB. FM-464 ($\lambda$ex/$\lambda$em = 515/640 nm) is a hydrophobic dye added to the media at a concentration of 2 $\mu$M to label the oil. The bacteria constitutively express EGFP ($\lambda$ex/$\lambda$em = 488/509 nm). The co-localization of magenta and green indicates that the dendritic branches are bacteria-stabilized tubes containing oil. (B) Confocal image of a bacteria-covered oil tube with the orientation director field overlaid. The red line in the director field is the central axis of the tube. (C) Dot product calculated for (B) between the cell axis and the tube axis (red) averaged over all cells in bin widths of 1.45 $\mu$m. (D) Image sequence showing the evolution of representative tubes. Due to rotation of the mother droplet, the location of the two protrusions, p1 and p2, respectively, rotate clockwise in subsequent frames.

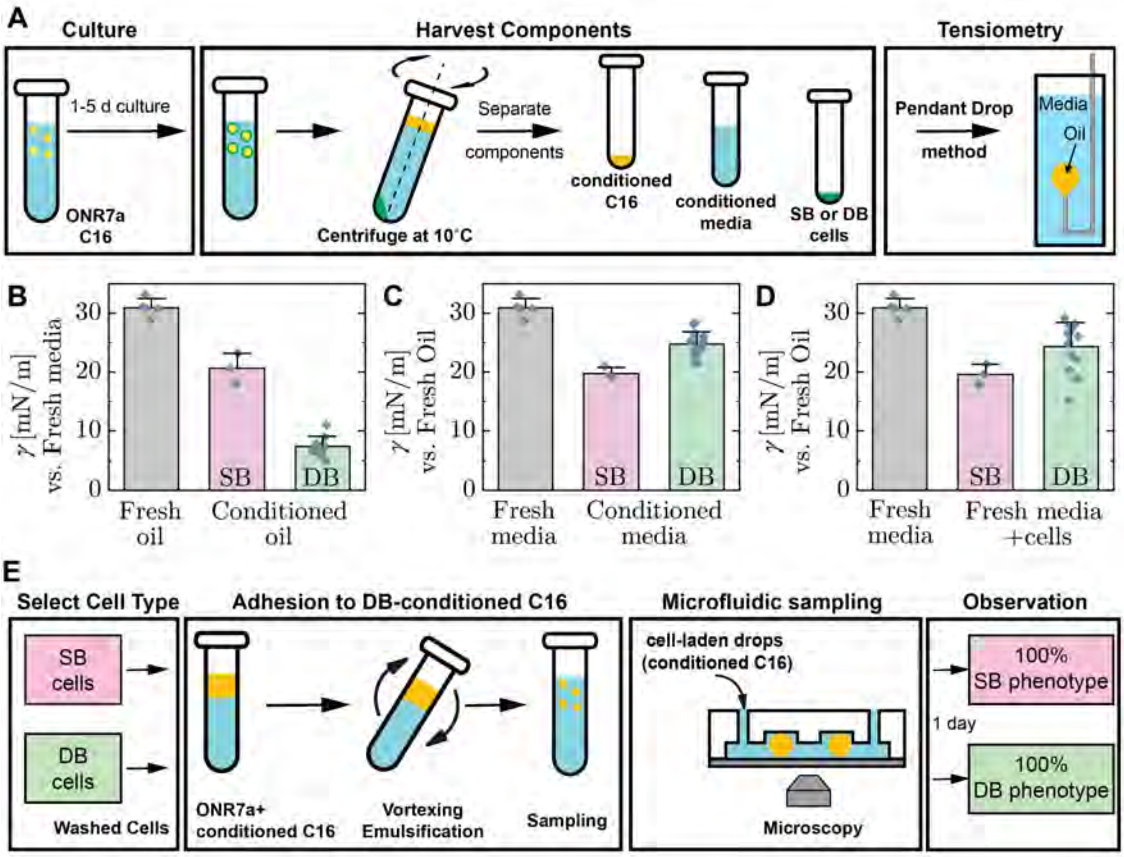

**Figure S6: Component separation and interfacial tension measurement using pendant drop method.** (A) Schematic showing the method of component harvesting and pendant drop method. See Methods for the detailed protocol. We measure the interfacial tension,$\gamma$, using the pendant drop tensiometry for the three fractions and plot the plateau values of $\gamma$. (B) $\gamma$ for conditioned C16 with fresh media ($n_{\text{control}}$=5; $n_{\text{SB}}$=3; $n_{\text{DB}}$=10; each measurement was done independently). This data is shown in **Fig. 2C** but is reproduced here to facilitate comparison. (C) $\gamma$ for conditioned media with fresh C16 ($n_{\text{control}}$=5; $n_{\text{SB}}$=3; $n_{\text{DB}}$=14; each measurement was done independently). The conditioned media was filtered through a 0.22 $\mu$m filter to remove all cells. (D) $\gamma$ for fresh media containing harvested cells that were washed (3x) after 1 day of culture with fresh C16 ($n_{\text{control}}$=5; $n_{\text{SB}}$=3; $n_{\text{DB}}$=14; each measurement was done independently). (E) Schematic describing the method we used to measure the phenotypic outcome from microfluidic sampling for pre-conditioned oil. We obtain the pre-conditioned oil from the 24 h liquid culture of DBs (see A). Here $\gamma \simeq 8$ mN/m (see B). We find that the observed phenotype remains constant despite the lowered $\gamma$ ($n_{\text{SB}}$= 2; $n_{\text{DB}}$=1).

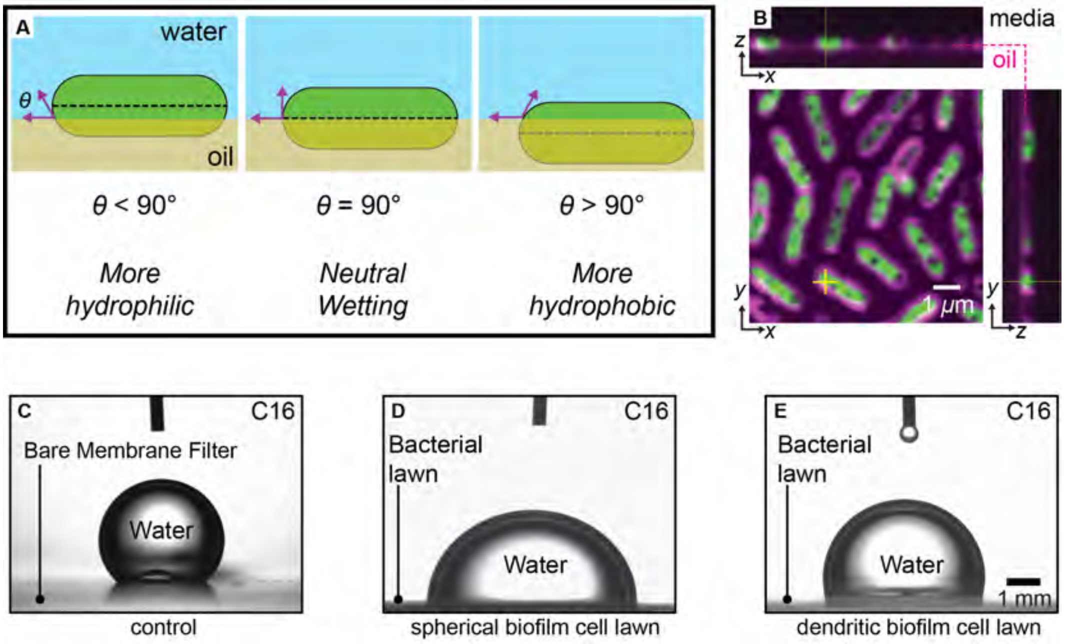

**Figure S7: C16 wettability of the cell surface.** (A) Schematic showing how spherocylinders of different surface-hydrophobicity will sit at the oil-water interface at equilibrium. (B) Confocal image of GFP-expressing DB cells at a flat oil-water interface that have been labeled with FM-464 (magenta). In addition to the cell membrane, FM-464 also stains the oil and can be used to visualize the water-oil interface (see faint magenta line in the orthogonal planes). The cells appear to have near neutral wetting. See Microfluidics Methods for experimental details. (C-E) Representative images of a 3-phase contact angle test for a water droplet submerged in C16 in contact with: (C) the bare membrane filter, where $\theta$=133 $\pm$ 5°; (D) the filter membrane supporting a bacterial lawn consisting of SB cells, where $\theta$=80 $\pm$ 8°; and (E) the filter membrane supporting a bacterial lawn of DB cells, where $\theta$=110 $\pm$10°. See Interfacial Properties Methods for experimental details.

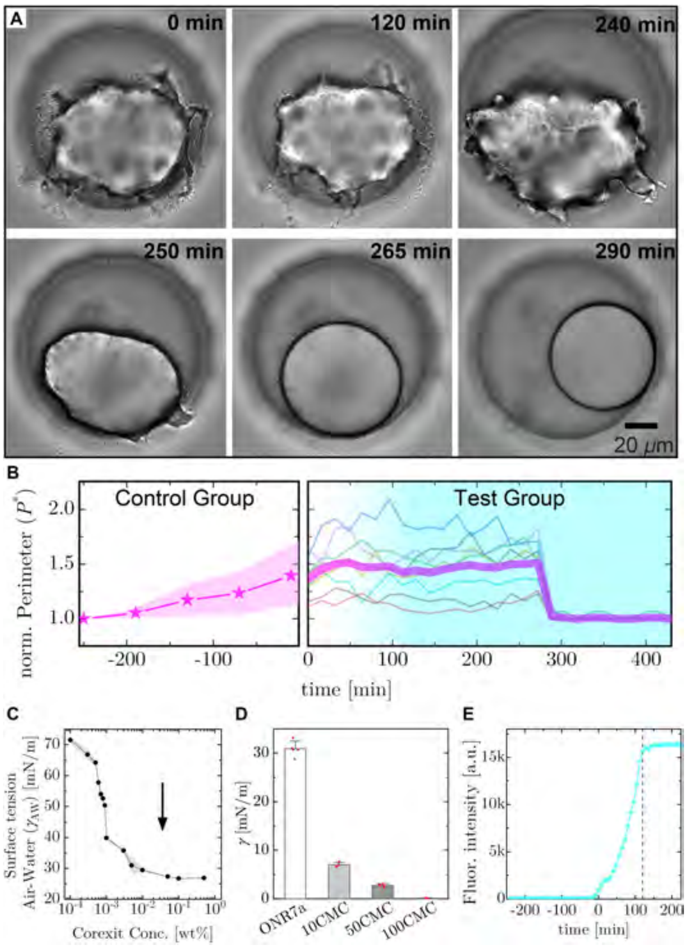

**Figure S8: Surfactant mixture disrupts DBs on oil drops.** (A) Time-lapse sequence showing the effect of a surfactant mixture, similar in composition to Corexit, on a typical DB growing on an oil droplet. We infuse media containing $50\times$ the critical micelle concentration (CMC) into the device, taking ∼250 min to reach the chamber. Here, 0 min indicates the time when the surfactants initially reach the chamber. These times should not be confused with $t_0$ in **Fig. 1**, which refers to the time of confluency on the droplets. (B) The normalized perimeter ($P^*$), measured at the drop equator, as a function of time under. (left) Evolution of $P^* = P/P_0$ for drops not exposed to surfactants; here, $P_0$ is the initial perimeter. The solid line and filled area represent the mean ± s.d. (n=15). To synchronize the time axis with the influx of surfactants into the device, we plot the control group from ∼ -250 min. (right) Evolution of $P^* = P/P_f$ for the test group, where $P_f$ is the final perimeter. At ∼280 min, biofilms abruptly detach and the drops become spherical. The thick magenta line represents the mean value, while the lines represent different droplets (n = 8). As in (A), 0 min refers to the moment when the surfactants first reach the chamber. Fluctuations in individual $P^*$ curves arise due to rotation of the droplets in traps. The cyan gradient shading indicates the surfactant concentration at the outlet of the drop chamber (see **Fig. S2A**), and reaches 95% of the steady-state concentration at ∼120 min, which we determine by calibrating the device. To generate DB, we culture the test group for ∼240 min prior to 0 min. (C) The air/water surface tension ($\gamma_{\mathrm{AW}}$) as a function of concentration of our mock Corexit dispersant mixture (See Methods for composition). We estimate the CMC to be $10^{-3}$ wt% and the arrow indicates 50CMC. The solid line and filled area represent the mean ± s.d. (n=3). (D) Oil-water interfacial tension between ONR7a media containing different concentrations of mock Corexit and C16. The bar labeled ONR7a is the control test. We normalize concentrations by the CMC, which was determined in (C). The error bars represent ± s.d. (n $\geq 3$). (E) Calibration curve for the flow conditions in this experiment, from which we estimate the concentration of the dispersants as a function of time. Fluorescence intensity is recorded as function of time at the chamber outlet using 3 $\mu$M FITC dissolved in the media containing mock Corexit (50CMC) infused at 0.5 $\mu$L/min. The solutes take ∼240 min to travel from the syringe to the chamber and requires an additional ∼115 min to reach 95% of the steady state concentration, indicated by the vertical dashed line. For this test, we label 0 min as the moment fluorescence is detectable at the chamber outlet.

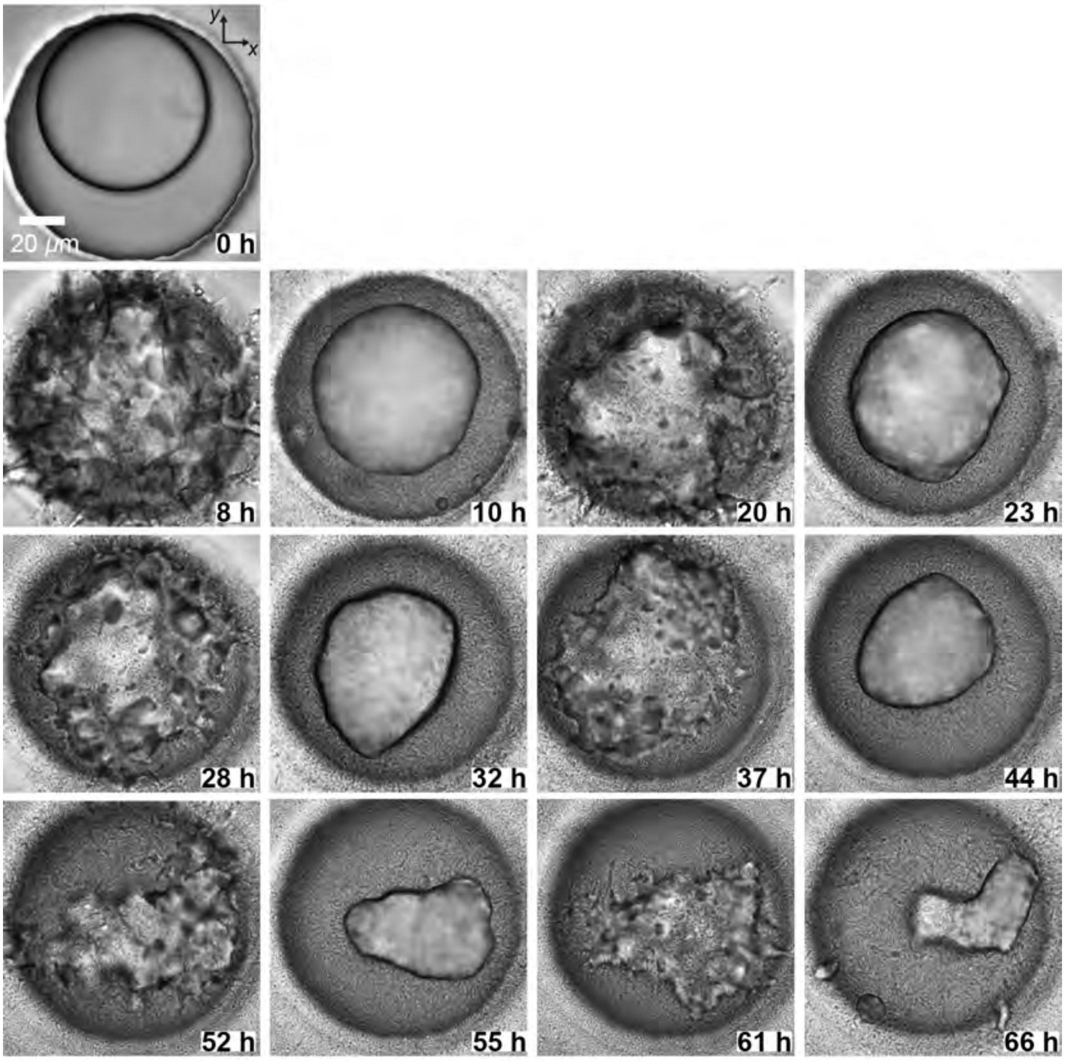

**Figure S9:** Oscillatory phase on a representative droplet, corresponding to **Supplementary Movie S5**. A cell monolayer is formed at 0h. Over the course of the next 65h, the biofilm phenotype oscillates between the SB and DB phases. The time-points at 10h, 23h, 32h, 44h, 55h, 66h (resp. 8h, 20h, 28h, 37h, 52h, 61h) are chosen as local minima (resp. maxima) of the oil/water interface length.

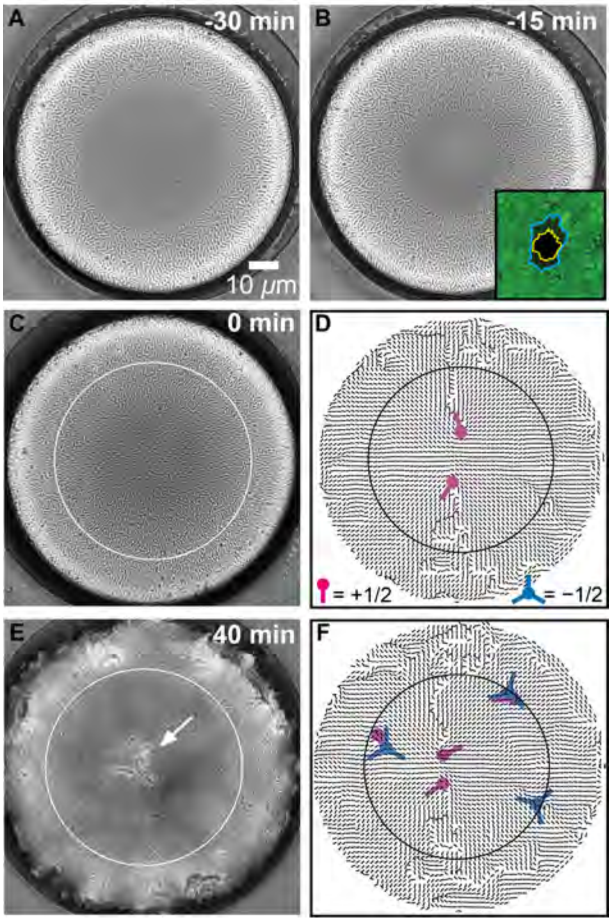

**Figure S10:** Flattened droplets develop two closely spaced $+1/2$ topological defects at $t_0$. All brightfield images are extended focus images taken at the bottom of the flattened droplet shown in **Fig. 4**. Acquisition times are indicated. (A,B) Interfacial tension of the droplet excludes cells from a circular region at the bottom of the droplet. (B,inset) Confocal image showing the excluded region at the bottom of the droplet, outlined in blue. Through this hole, the excluded region at the top of the drop, outlined in yellow, is visible. (C) Pressure in the cell layer due to division drives cells into the excluded region, generating a monolayer. The circle indicates the flat region of the droplet due to contact with the underlying cover glass. (D) The director field for the image in (C) and two closely spaced $+1/2$ defects (magenta) are shown. (insets) Magenta and blue icons indicate topological defects of charge $+1/2$ and -1/2, respectively. (E) A dimple forms at the center of the flat region, indicated by the arrow. (F) The director field for the image in (E) with $+1/2$ defects (magenta) and -1/2 defects (blue).

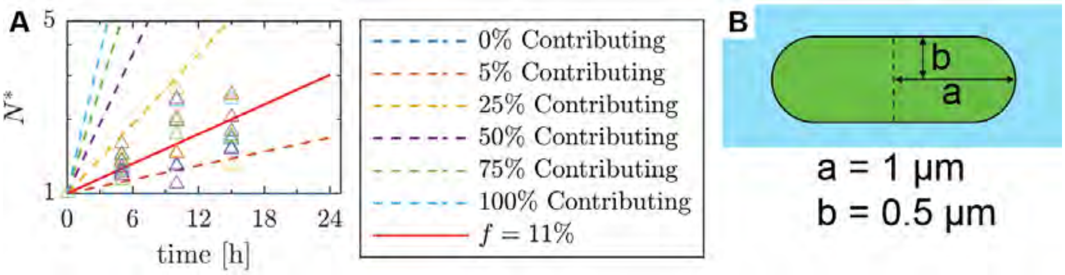

**Figure S11:** Estimated number of cells at the interface that cause an increase in surface area. (A) We plot $N^*$ for DBs as a function of time ($\Delta$). $N^*$ is calculated from the data of 10 different drops shown in **Fig. 1D** using $N^* = \frac{\phi}{s_c} S^*$, which comes from **Eq. 9**. Here, $s_c$ is the cross-sectional area of a bacterium (1.8 $\mu m^2$) and $\phi$ is the packing fraction (0.65). The initial value of $N^*$=1 represents the number of cells in a monolayer. The solid line is the best fit to the data by fitting the predicted number of cells that reside at the droplet surface assuming that only a percentage of cells of all cells participate in expanding the interface from **Eq. 8**. The dashed lines show the predicted number of cells that would reside at the interface, assuming that only a percentage $f$ of all cells participate to expand the interface. These cells may be lost from the interface during each division cycle and thus are not active participants in the interface. (B) Typical dimensions of a *A. borkumensis* cell when cultured using C16 as the carbon source.

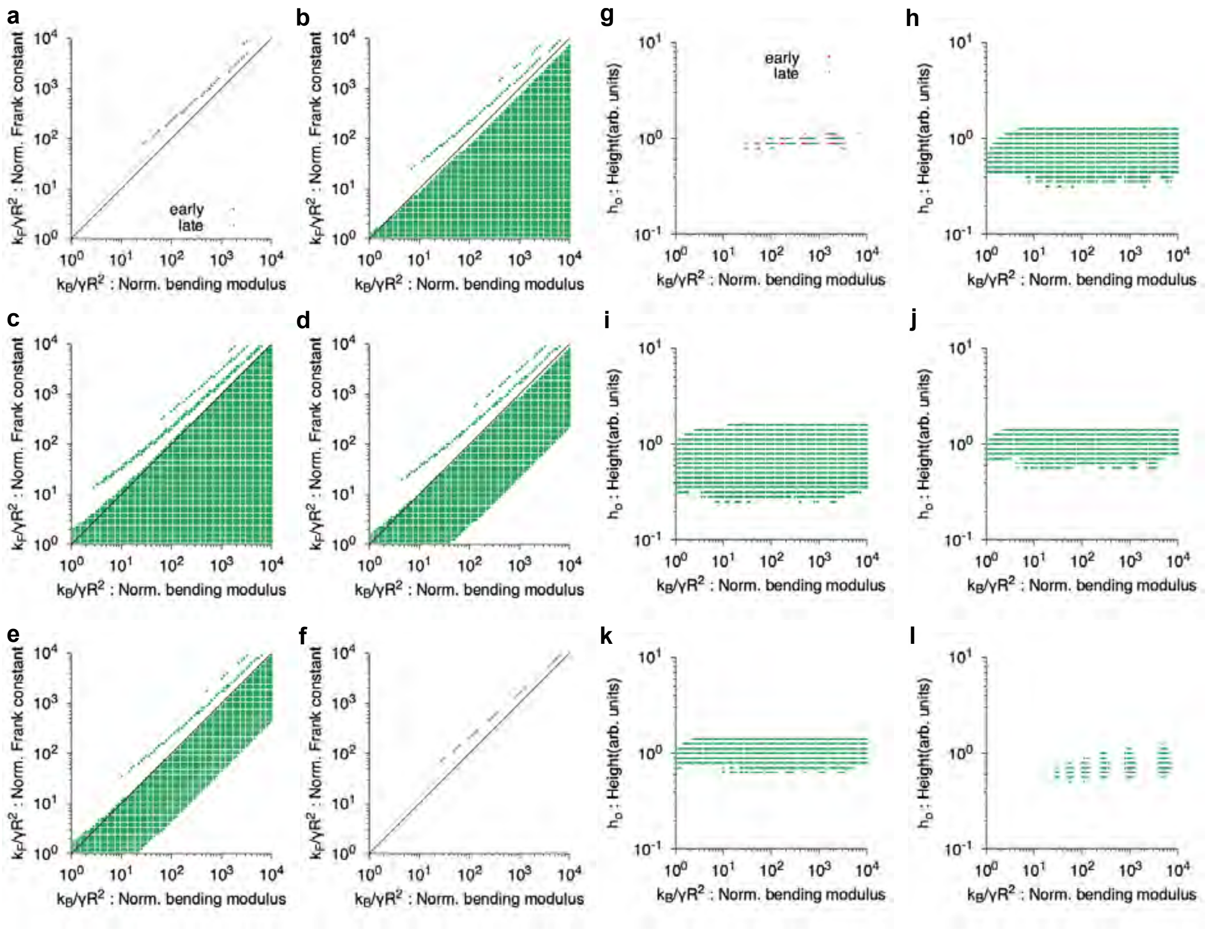

**Figure S12:** Region of the parameter space whereby the error function $\mathcal{E}$ given by Eq. (76) is at most $20\%$ larger than the absolute minimum error $\mathcal{E}_{min}$. Panels **a-f** show a cross-section of the parameter space on the plane $\kappa_B/\gamma R^2$-$\kappa_F/\gamma R^2$, and panels **g-l** on $\kappa_B/\gamma R^2$-$h_0$. The experimental cases are ordered from a-f and g-l as in Table S2 and $R$ corresponds to $R_{\mathrm{dimple}}$.

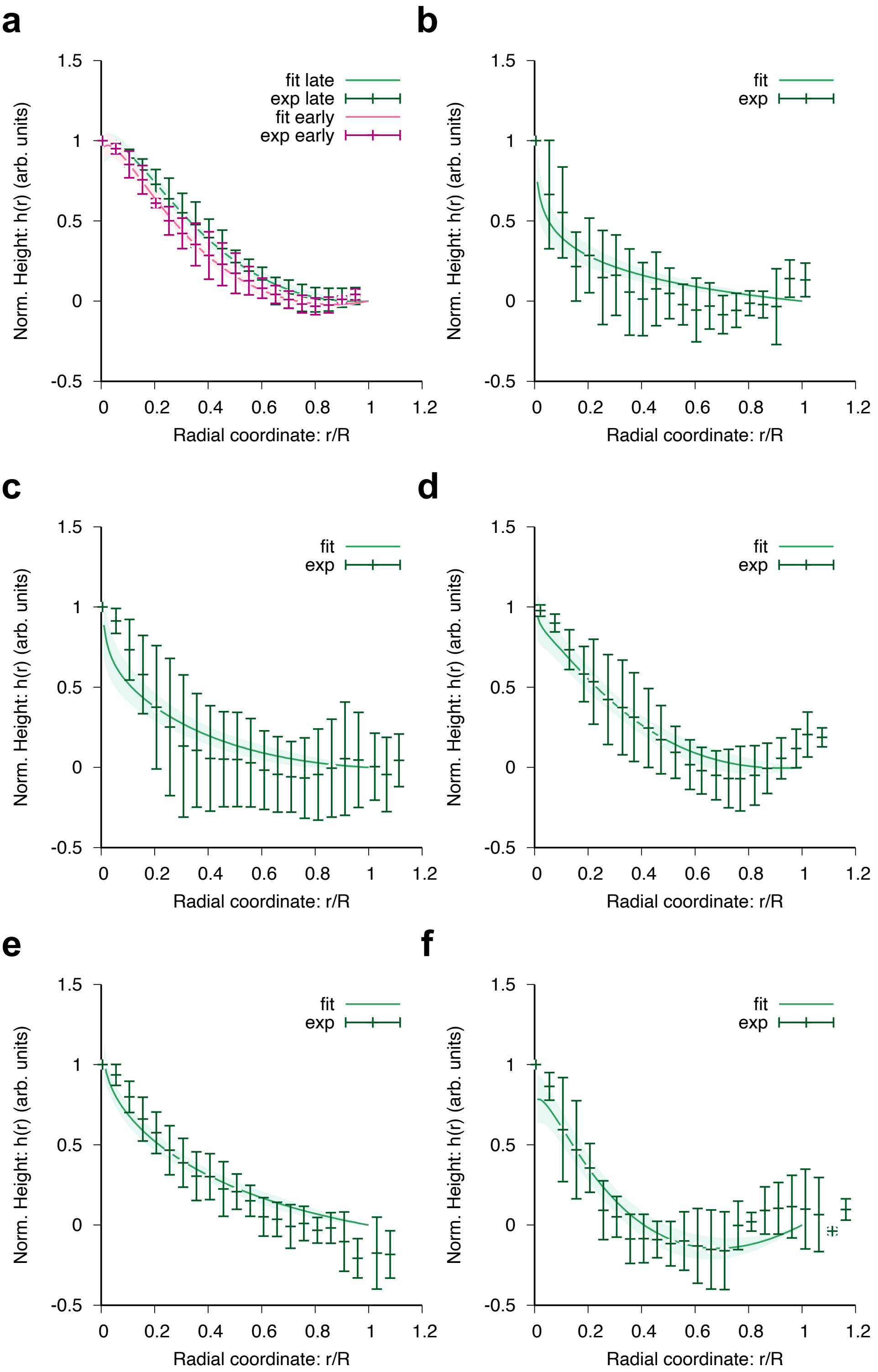

**Figure S13:** Theoretical fits to experimental data. (a-f) Dimple profiles as a function of the radial coordinate $r/R$. In (a-f), the green lines are theoretical fits while the dark green points are experimental profiles ($n = 4$). In (a), the magenta line is a theoretical fit while the dark magenta points are experimental profiles ($n = 4$). Error bars in theoretical fits, which are represented as shaded areas, correspond to the standard deviation of parameter values that lead to $\mathcal{E} < 1.2\mathcal{E}_{min}$ and are then averaged in time. In the experimental curves, the error bars correspond to the standard deviation for $n = 4$ that are averaged in time. In experiments, the averaged dimple radius ranges from $R = 25 - 35$ $\mu$m, while the averaged height ranges from $h = 4 - 8$ $\mu$m. The experimental cases are ordered from (a-f) as in Table S2 and $R$ corresponds to $R_{\text{dimple}}$

.

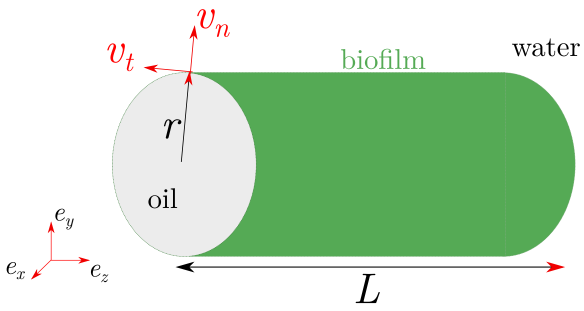

**Figure S14:** Sketch of the model dendrite geometry: a tube of constant radius $r$, base-to-tip length $L$, ended by a half-spherical cap of radius $r$.

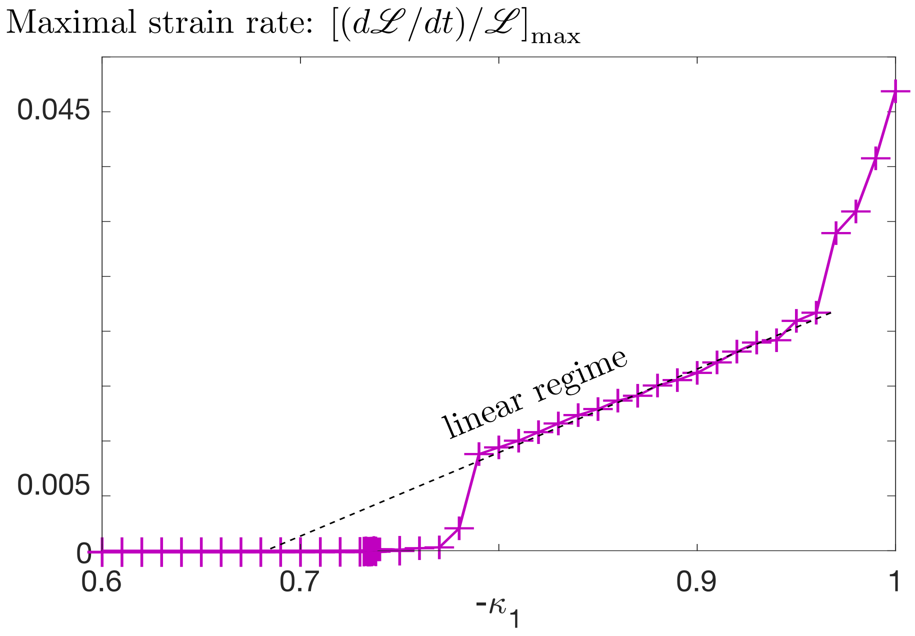

**Figure S15:** Phase-field model with no coupling to the bacteria density: maximal strain-rate (defined as $\dot{\mathscr{L}}/\mathscr{L}$, where $\mathscr{L}$ is the interface length, Eq. 62) as a function of a fixed effective surface tension parameter $\kappa_1$ (linear scale).

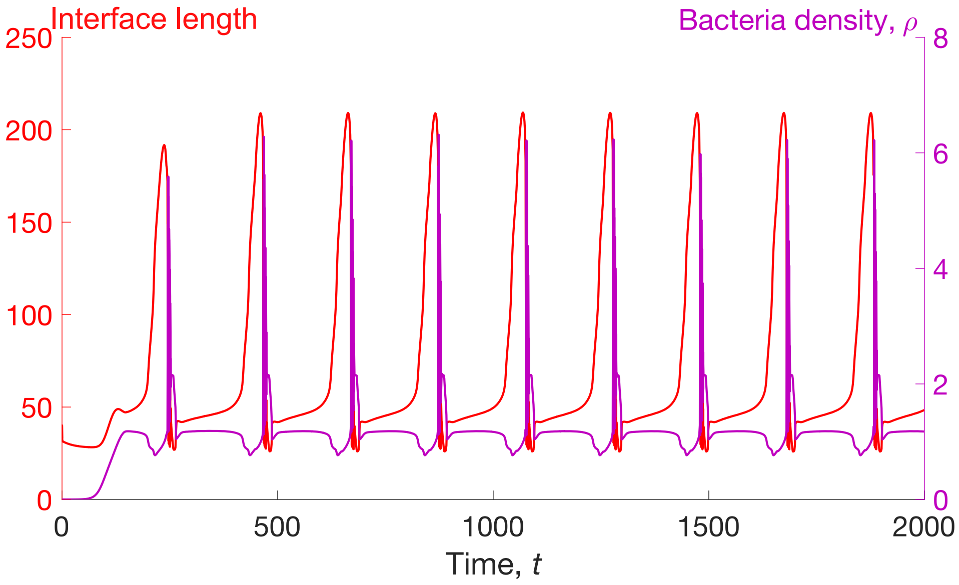

**Figure S16:** Phase-field model with coupling to the bacteria density. Time evolution of the total interface length $\mathscr{L}$ (red, left axis) and density $\rho$ (magenta, right axis) for the oscillatory regime parameter set, $k_1 = 5.4, \rho_H = 1.2$, with all other parameters fixed in Sec. S7.5 **Supplementary Movie 4**. During the rapid tube expansion phase, the density is lower than $\rho_H = 1.2$.

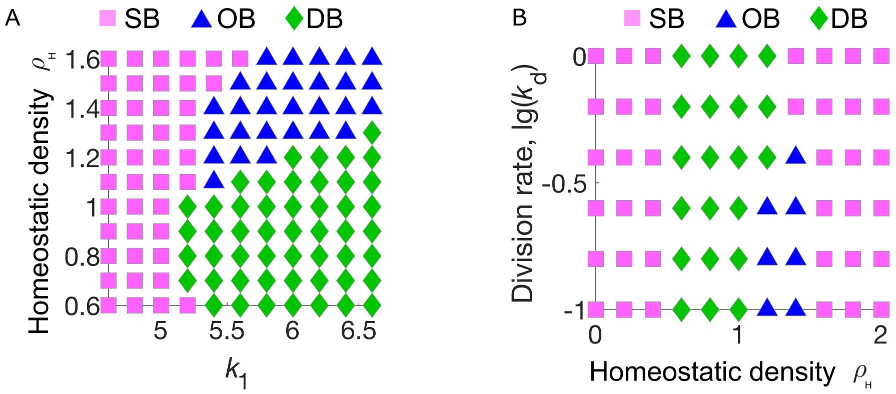

**Figure S17:** Phase-field model with coupling to the bacteria density. (A-B) Phase diagrams of the modeled spherical biofilm (magenta squares), oscillatory biofilm (blue triangles) and dendritic biofilm (green diamonds) phenotypes as a function of model parameters (A) $\rho_H$ and $k_1$ and (B) $k_1$ and $\rho_H$.

### S10 Supplementary Tables

| | Interfacial Tension $\gamma$ (mN/m) of respective fractions | | |
|---|---|---|---|
| | Cond. oil vs Fresh media | Cond. media vs. Fresh oil | Cells (in Fresh media) vs. fresh oil |
| SB Phenotype | 20.6 ± 2.1 | 19.8 ± 0.8 | 19.6 ± 1.4 |
| DB Phenotype | 7.4 ± 1.6 | 24.7 ± 2.0 | 24.3 ± 3.9 |

**Table S1:** Interfacial tensions of the different phases. The control tests (fresh C16 versus fresh ONR7a) has an $\gamma$ = 30.9 ± 1.4 mN/m.

| Parameter | #1 Early | #1 Late | #2 | #3 | #4 | #5 | #6 |
|---|---|---|---|---|---|---|---|
| $\kappa_F/\kappa_B$(dimensionless) | $1.8 \pm 0.2$ | $2.0 \pm 0.4$ | $< 4$ | $< 5.5$ | $< 5$ | $< 4$ | $1.7 \pm 0.3$ |
| $h_0$(arb. units) | $1.0 \pm 0.1$ | $0.9 \pm 0.1$ | $0.8 \pm 0.2$ | $1.0 \pm 0.3$ | $1.1 \pm 0.2$ | $1.1 \pm 0.2$ | $0.8 \pm 0.2$ |

**Table S2:** Estimation of material parameters for a membrane with nematic order. The parameter $\kappa_F/\kappa_B$ corresponds to the ratio between the energetic cost due to distortion of the director field and due to bending deformations. The error bars correspond to the standard deviation in the region of the parameter space that satisfies $\mathcal{E} < 1.2 \times \mathcal{E}_{min}$. 'Early' corresponds to the green curves while 'late' corresponds to magenta curves shown in Fig. S13. 'Early' corresponds to height profiles at $t = 20$ min post-confluency while 'late' is at $t = 26 \pm 3$ min post-confluency.

#### S11 Supplementary Movies

**Movie S1:** Spherical biofilm (SB) phenotype. Full time-lapse sequence corresponding to **Fig. S3B**.

**Movie S2:** Dendritic biofilm (DB) phenotype. Full time-lapse sequence corresponding to **Fig. S3C**.

**Movie S3:** Effect of a surfactant on a dendritic biofilm (DB). Full time-lapse sequence corresponding to **Fig. S8**.

**Movie S4:** Oscillatory biofilm (OB) phenotype. Full time-lapse sequence corresponding to **Fig. 3G** in the main text and to **Fig. S9**.

**Movie S5:** Phase field simulations in the (left) spherical biofilm regime, $k_1 = 5.0, \rho_H = 1.2$; (middle) oscillatory biofilm regime, $k_1 = 5.4, \rho_H = 1.2$; (right) stable dendritic biofilm regime, $k_1 = 5.4$, $\rho_H = 1.0$, with all other parameters fixed in Sec. S7.5.

**Movie S6:** Bacterial invasion of the flat regions at the bottom of a squeezed droplet followed by biofilm-mediated droplet dimpling. Full time-lapse sequence corresponding to **Fig. 4** in the main text and **Fig. S10**.

Version : 17/10/2022 at 08:48:13 (UTC)
